# Directing hierarchical cell fate decisions through sequential pulses of minimal signaling alphabets

**DOI:** 10.64898/2026.07.28.741174

**Authors:** Yufei Ji, Zhiyuan Li

## Abstract

Inductive signals can direct cell fate decisions, yet the diversity of cell identities far exceeds the available signaling repertoire. While developing systems resolve this paradox by applying temporal sequences of minimalistic signals to hierarchically organized gene regulatory networks (GRNs), the rules governing robust sequential fate navigation remain largely unknown. To address this, we modeled how sequential signals steer trajectories toward arbitrary fates within a Waddington-like manifold, enumerating parameters and combinatorial logics of cross-inhibition and self-activation (CIS) modules, and various ways of applying the sequential signals. We uncover a fundamental conflict between commitment stability and inductive plasticity that severely limits sequential fate navigation. Crucially, introducing non-inductive gap intervals systematically resolves this bottleneck by dissipating kinetic leakage without sacrificing upstream stability. Finally, we demonstrate that polarization-division cycle can autonomously generate signals meeting these temporal requirements. Together, our findings provide a minimal model demonstrating the feasibility and design principles for driving hierarchical cell fate decisions through sequential pulses of a restricted signaling repertoire.

## Introduction

Waddington’s epigenetic landscape has served as an important conceptual paradigm for cell fate decisions for over six decades[1, 2]; yet, many questions remain in bridging this intuitive concept with quantitative theories[3, 4]. Specifically, while the landscape elegantly captures the spontaneous differentiation of a cell from the stem cell state, the precise controllability of its terminal state is unresolved: as a cell navigates the landscape from pluripotency to terminal differentiation, how is its trajectory reliably induced into a targeted “valley”? This directed controllability is crucial for embryogenesis and regenerative medicine[5], where the precise spatio-temporal patterning of distinct cell types organizes tissue function. Conventionally, targeted fate decisions during lineage branchings are dictated by inductive signals[6, 7]. For example, in the mammalian pre-implantation embryo, the next fate of the inner cell mass (ICM) is dictated by fibroblast growth factor 4 (FGF4): ICM cells exposed to high FGF4 canalize into the primitive endoderm (PrE), whereas those experiencing low signaling maintain the epiblast (Epi) trajectory[8, 9]. Zooming into any isolated developmental branching point, one can almost invariably identify a local, signal-induced decision.

However, scaling this intuitive picture to the global organismal level exposes the “Toolkit Paradox”[10, 11]: biologically documented developmental signaling pathways belong to a strictly conserved alphabet of merely 7 to 12 core families (e.g., Wnt, TGF-*β*, FGF, Hedgehog)[12], which stands in stark contrast to the several hundred distinct cell types comprising a vertebrate body. If inductive signals alone dictated the choice of branching directions at each fork, the number of reachable terminal states would be severely upper-bounded. Formulating the cell fate decision as a dynamical mapping,

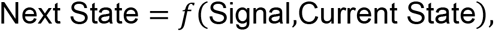

this dimensional mismatch suggests that the current state must act jointly with the inductive signals. Returning to the landscape metaphor: under an idealized scenario with only a two-signal alphabet (sA and sP, representing “anterior” and “posterior”), the system can theoretically access an unlimited number of valleys, provided that the cell’s current position on the landscape determines the local action of the signal. For example, the exact same signal sP pushes a cell into the second valley *AP* if it currently resides at state *A*, but into the fourth valley *PP* if it resides at state *P* [13, 14]. Consequently, high-dimensional fate induction under a minimal signaling alphabet mandates historical memory: the accumulated developmental trajectory determines the current state[15], thereby contextually rewiring the cell’s response to future stimuli (Figure 1A).

**Figure 1:**
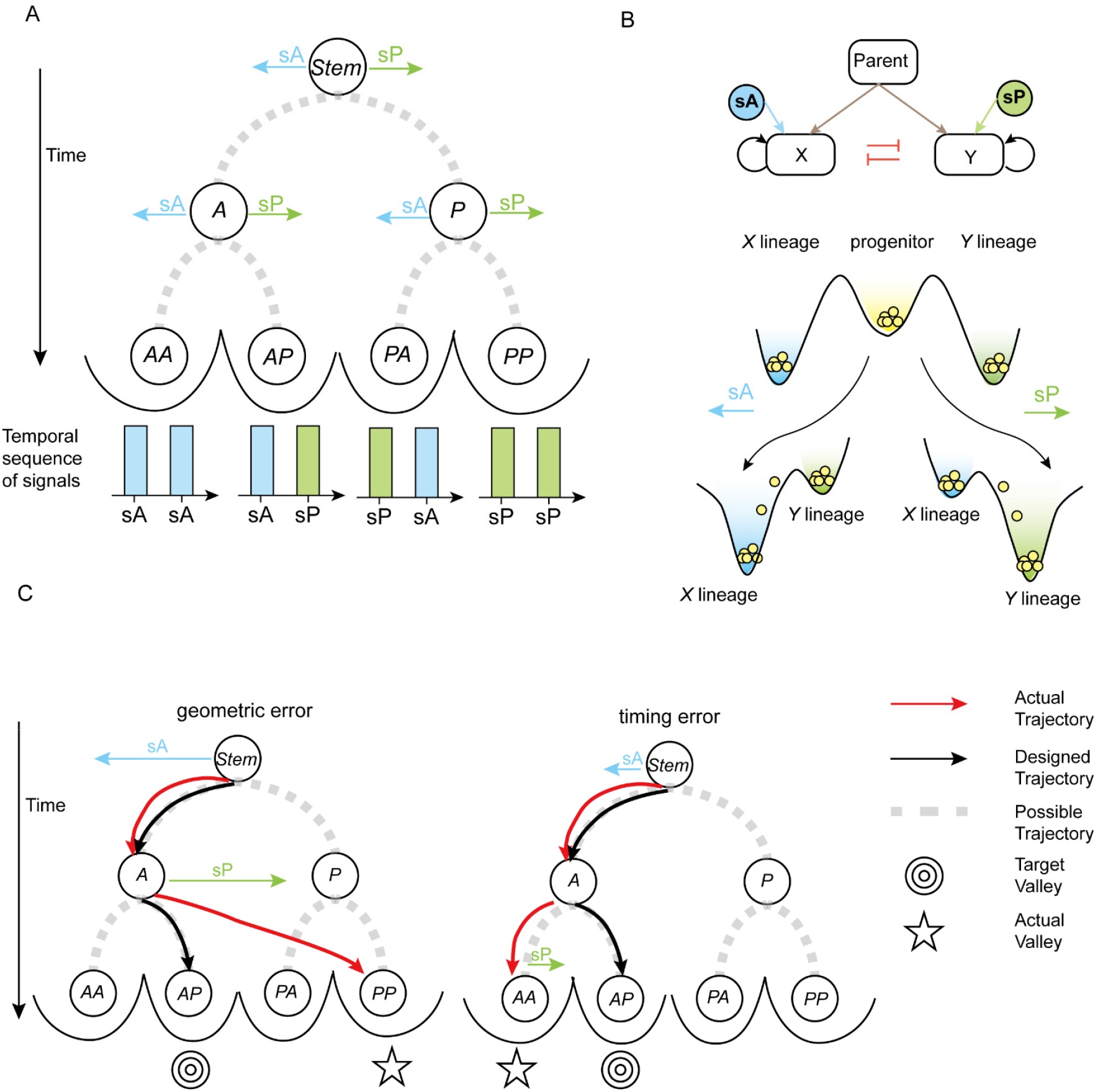
Sequential cell fate induction in hierarchical landscapes and its dynamic vulnerabilities. (A) Schematic of the hierarchical induction. Gray dashed lines represent all possible trajectories. Blue and green arrows represent external inductive signals, sA and sP, respectively. Applying specific temporal sequences of these signals drives the cell into distinct valleys (e.g., using the signal sequence sA -> sA to induce the *AA* fate). (B) Schematic of CIS module with a parent node and corresponding landscapes. Top: Schematic of CIS module with a parent node. Black and red arrows represent self-activation and cross-inhibition, respectively. Blue and Green arrows represent external inductive signals, sA and sP, respectively. Brown arrows represent parent activation. Bottom : Signal-induced landscape deformation. Yellow, blue and green valleys represent the uncommitted progenitor, *X* lineage and *Y* lineage, respectively. Inductive signals tilt the landscape to canalize the progenitor into the corresponding differentiated valley. (C) Schematic of geometric error and timing error. Gray dashed lines represent all possible trajectories. Black and red arrows represent designed trajectories and actual trajectories, respectively. Blue and green arrows represent external inductive signals, sA and sP, respectively. Left panel: Schematic of geometric error. The signal-induced landscape deformation is so extreme that the subsequent sP overwrites the previously established *A* fate, driving the cell into the unintended *PP* valley, rather than the target *AP* valley. Right panel: Schematic of timing error. the temporal switch from sA to sP occurs late. Consequently, the cell irreversibly falls into *AA* valley before sP arrives.

Mechanistically, multicellular organisms resolve this constraint by embedding state-dependence within hierarchical gene regulatory topologies. A pervasive, conserved motif in such hierarchies is the cross-inhibition and self-activation (CIS) module [16, 17]. Driven by two antagonistic master transcription factors (denoted as *X* and *Y*) that self-activate and cross-inhibit, a single CIS circuit can generate a classic cell-fate landscape comprising two terminal differentiated attractors (characterized by mutually exclusive X^high^/ Y^low^and X^low^/Y^high^ expression profiles) and a central attractor representing the uncommitted progenitor (with low expression of both factors). External inductive signals selectively targeting either X or Y deform this landscape, driving the progenitor to canalize into the corresponding differentiated valley (Figure 1B). In a developmental hierarchy, multiple CIS modules are wired in series: the entry of a cell into the differentiated valley *X* unlocks the accessibility of a downstream CIS module, rendering it competent to respond to the next wave of signals; simultaneously, all regulatory modules downstream of the unchosen fate *Y* remain silent and unresponsive to signals[18]. For instance, during early mammalian embryogenesis, the primary CIS of Oct4 and Cdx2 segregates the inner cell mass (ICM, Oct4^+^) from the trophectoderm (Cdx2^+^) [19]. The winning factor Oct4 subsequently primes the downstream Nanog–Gata6 CIS circuit exclusively within the ICM, rendering these cells competent to respond to the aforementioned FGF4 signal [20]; conversely, the trophectoderm branch lacks Oct4 and remains insulated from FGF4-mediated Epiblast induction. Through this hierarchical state-dependence, a compact set of signals, when deployed in precise temporal sequences (e.g., sA->sA->sP->sP->sA), can theoretically unroll an unbounded fate-branching tree [21].

Mathematical modeling has been widely applied to conceptualize the attractor landscapes of cell fate decisions [3, 4, 22]. In theoretical biology, minimal proof-of-concept models are often most valuable not for achieving a detailed simulation of reality, but for identifying the critical assumptions that separate a functional mechanism from a failing one. While previous models have elegantly mapped the steady-state topography of developmental landscapes [4, 23] or the local dynamics of isolated branching events [24], they largely focus on the static shape of the valleys. However, in the idealized two-signal hierarchy described above, the system has only three effective topographies: one governed by signal sA, another by signal sP, and a third representing the state where neither signal is present. Basically, applying a temporal sequence of signals (e.g., sA->sP) corresponds to switching the underlying landscape back and forth between the three phase-spaces, dynamically “shaking” the cell state toward its desired target valley. This suggests that sequential fate induction is inherently a dynamic non-equilibrium process rather than a static equilibrium problem.

Intuitively, steering a system through time-varying transients is far more challenging than navigating a static landscape. Consider the sequential induction of the *AP* fate via the signal sequence sA->sP. Because inductive signals are broadcast globally to all genes in the network, targeted induction is vulnerable to two types of failures. First, geometric errors can occur if the signal-induced landscape deformation is insufficient or overly extreme. For instance, if the subsequent sP signal induces an excessive deformation of the global landscape towards the *P* state, it can overwrite the previously established “sA” history, driving the cell into the unintended *PP* valley (Figure 1C, left panel). Second, even if the landscape deformation is perfectly tuned, timing errors can destroy the trajectory. If the temporal switch from sA to sP occurs too late, the cell state may have already irreversibly crossed the downstream bifurcation threshold, causing it to fall uncontrollably into the *AA* valley despite the arrival of the correct signal sP (Figure 1C, right panel). Therefore, the critical kinetic bottlenecks and theoretical limits of this dynamic, multi-step induction process remain to be fully explored.

In this study, we seek to identify the essential regulatory logical and signaling temporal constraints for reliable hierarchical fate induction by sequentially reusing two types of signals only. We first systematically enumerate all 52 Boolean models within a hierarchical CIS framework[25], and reveal a set of conserved state-transition rules and attractor basin structure that serve as the mechanistic foundation for preventing fate reversal and premature activation. Next, in continuous ODE model, we further identify an optimal signal switching period 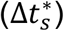 [26]and demonstrate that non-inductive gap intervals (Δ*t*_*b*_)[27] are necessary and sufficient for robust induction at arbitrary depths under realistic kinetic constraints. Finally, we show that these control principles can be autonomously implemented through self-organized polarization-division cycles, providing a concrete biological realization for hierarchical fate specification.

## Results

### Enumeration of Boolean models reveals a conserved state-transition structure for successful hierarchical induction

We first ask whether Boolean models can perform sequential fate navigation, i.e., achieving targeted fate by sequentially reusing a two-signal alphabet (Figure 2A). To formalize our conceptual framework, we construct an idealized gene regulatory network comprising a binary tree of interconnected CIS modules, as illustrated in Figure 2B. Let Σ = {A, P} denote a binary alphabet associated with the two globally broadcast inductive signals, sA and sP. Every individual transcription factor (TF) in the network is assigned a unique hierarchical coordinate represented by a string *s* ∈ *Σ*^*k*^ of lineage depth *K* ≥ 1 (e.g., TF A and TF P at depth *K* = 1; TF AA at depth *K* = 2).

**Figure 2.**
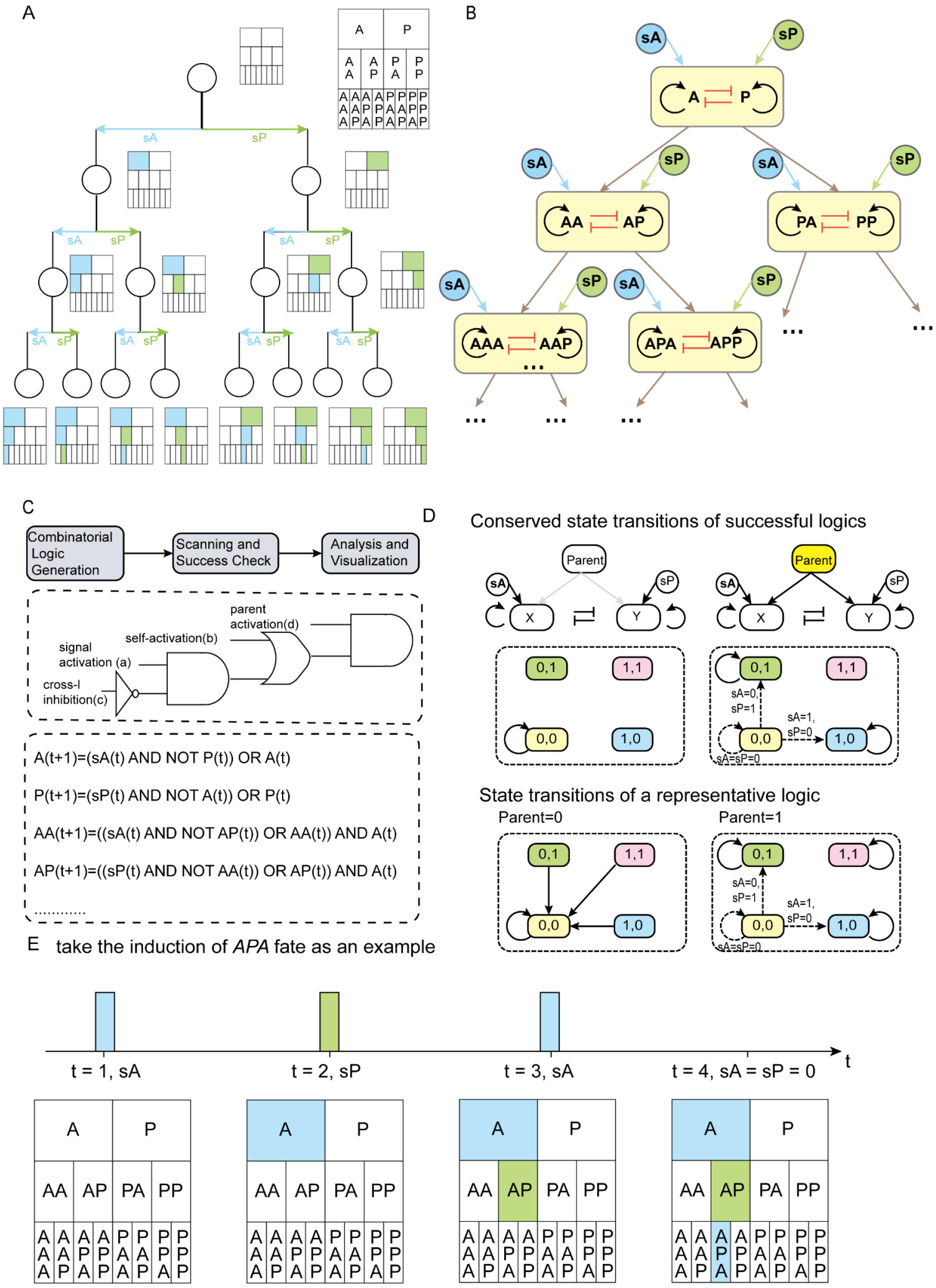
Hierarchical network topology and conserved Boolean logic for targeted sequential fate induction. (A) Schematic of desired hierarchical induction. Circles denote individual cells across developmental stages; corresponding block diagrams represent their internal transcription factor (TF) state vectors. Sibling branches bifurcate under the mutually exclusive action of globally broadcast signals sA (blue) and sP (green). The terminal state vector should record the cumulative history of inductive pulses (e.g., the signal sequence sA -> sP -> sA uniquely canalizes the cell into the *APA* terminal basin). (B) Topology of an idealized gene regulatory network comprising a binary tree of interconnected CIS modules. Sibling TFs sharing a common parent prefix are organized into cross-inhibition and self-activation (CIS) modules (yellow boxes). Sibling nodes are driven by four core regulatory interactions: external signal activation (a, blue/green arrows), self-activation (b, black loops), cross-inhibition (c, red blunted lines), and parent activation (d, brown arrows) (C) Flowchart of the Boolean logic screening process. Top: Computational screening workflow across all 52 non-redundant logic models. Middle: The equivalent logic gate circuit governing the input integration of (a)-(d) for a representative successful model. Bottom: The explicit Boolean state-update equations derived from this representative logic. (D) Conserved state-transition structure for successful hierarchical induction (top panel) and the state-transition graph of the exemplary logic shown in (C) (bottom panel). Transitions are gated by the status of the parent node. Solid black arrows indicate transitions occur independent of inductive signals, and dashed black arrows represent signal-driven transitions. Background colors denote cell states: yellow for the dual-low uncommitted progenitor state, blue/green for the single-high differentiated states, and pink for a dual-high state. (E) Example of a successful targeted induction for the *APA* cell fate. Signals sA and sP are applied sequentially at time steps t = 1, 2, and 3, respectively. Active transcription factors (Boolean value = 1) are highlighted in color (blue for sA-activated, green for sP-activated nodes), while inactive nodes (Boolean value = 0) remain uncolored. Crucially, at t = 4, external signaling is totally withdrawn (sA = sP = 0).

Structurally, for any TF node denoted by *s* = *iσ* (where *i* ∈ *Σ*^k−1^ is the parent prefix string and *σ* ∈ *Σ*^l^ is the terminal symbol): The prefix *i* specifies its upstream parent TF node, and the terminal symbol *σ* determines which signal activates the TF node — it is activated by sA if *σ* = *A*, and by sP if *σ* = *P*. For example, the TF APA should be switched ON if and only if the cell has experienced the ordered signal sequence sA->sP->sA. Similarly, each terminal fate is named by the TF activated in the corresponding fate.

The global network architecture is governed by three recursive topological settings:

1. Intra-module CIS motif: A module consists of a pair of sibling TFs, iA and iP, sharing a common parent prefix string *i* ∈ *Σ*^∗^ (where *i* = *ϵ* denotes the empty root prefix). Within each module, TF iA and TF iP self-activate and cross-inhibit.
2. Hierarchical activation: The inter-module wiring is feedforward in a binary-tree fashion. The parent TF *i* activates its direct downstream module (iA, iP).
3. Global signal perception: The inductive signals sA and sP are broadcast globally across the entire network. Signal sA selectively targets all competent TFs ending in the symbol ‘A’ (i.e., TF A, PA), while sP targets all competent TFs ending in ‘P’ (TF iP), regardless of their hierarchical depth *k*.

Under these setting, each TF perceives four regulatory inputs: one external inductive signal (sA or sP) (a), two intra-module interactions, including self-activation (b) and cross-inhibition (c), and the cross-module parent activation (d). Even when strictly adhering to the three topological rules above and assuming that all modules share identical logic gates, the distinct Boolean combinations of inputs (a)-(d) still yield 52 unique models after eliminating network symmetries (Figure 2C).

To determine which combinatorial logics can successfully drive sequential differentiation toward bottom-layer target cell fates, we systematically evaluated the network dynamics of all 52 candidate models under sequential signal ordered trains of signals (e.g., applying sA->sP->sA sequentially to target the *APA* fate; Figure 2E). We define a “successful induction” as the fulfillment of two steady-state conditions post-induction: (i) lineage activation, where the targeted TF and its ancestral lineage are stably activated (1); and (ii) lateral suppression, where all non-target branch nodes remain strictly silent (0). This definition captures both lineage fidelity and exclusivity. Our screening revealed that the majority of the candidate logics failed to satisfy these criteria; out of the 52 enumerated models, only 4 successfully induced all target lineages, demonstrating that the capacity for hierarchical fate specification is a severely constrained logical property.

Despite their distinct Boolean algebraic expressions, all 4 successful combinatorial logics share common state transition structures. Examining the state-transition diagram for any pair of sibling TFs in a module (e.g., AA and AP; Figure 2D, Figure S1) reveals three conserved structural rules:

1. Parent-dependent gating: When the parent node is inactive (0), the uncommitted progenitor state (0,0) exclusively undergoes self-transitions, acting as an insulated sink regardless of the applied external signal.
2. Signal-dependent fate selection: Only when the parent node is active (1) and the system resides in the uncommitted progenitor state (0,0), does the applied signal dictate the state transition trajectory.
3. Commitment irreversibility: when the parent node is active (1), the established differentiated states, (0,1) and (1,0) exclusively undergo self-transitions, remaining permanently locked regardless of subsequent signal changes.

Collectively, these three rules enforce a two-stage developmental check: (i) undifferentiated cells become competent to respond to inductive signals only after their upstream lineage has been turned ON, and (ii) once a cell commits to a local basin, it rejects subsequent inductive signal overwriting. Together, these Boolean constraints define a conserved state-transition structure required to navigate a high-dimensional Waddington manifold using a reused, lower-dimensional signaling alphabet.

### Continuous ODE models confirm that conserved attractor basin structure underlies hierarchical induction in realistic dynamical regimes

While Boolean models capture the state-transition rules basic to hierarchical fate induction, they inherently abstract away the continuous kinetics that characterize real gene regulatory networks, specifically the finite response times and concentration-dependent gating. We therefore asked whether and how the conserved state-transition rules identified in the discrete Boolean model map onto conserved phase-space geometry in continuous dynamical systems. To address this, we extended the conceptual framework in Figure 2C into a deterministic ordinary differential equation (ODE) model. To model transcriptional regulation analytically, we replaced the discrete Boolean logic gates with continuous, steep sigmoidal approximations using Heaviside step functions (see Methods for details). Crucially, this continuous formulation naturally introduces two independent concentration thresholds: the intra-module interaction threshold (*Kintra*), governing the within-module self-activation and cross-inhibition, and the inter-module gating threshold (*Kinter*), determining whether a downstream transcription factor is unlocked (Figure 3A). This framework allows us to directly map discrete state-transition rules onto continuous expression manifolds.

**Figure 3.**
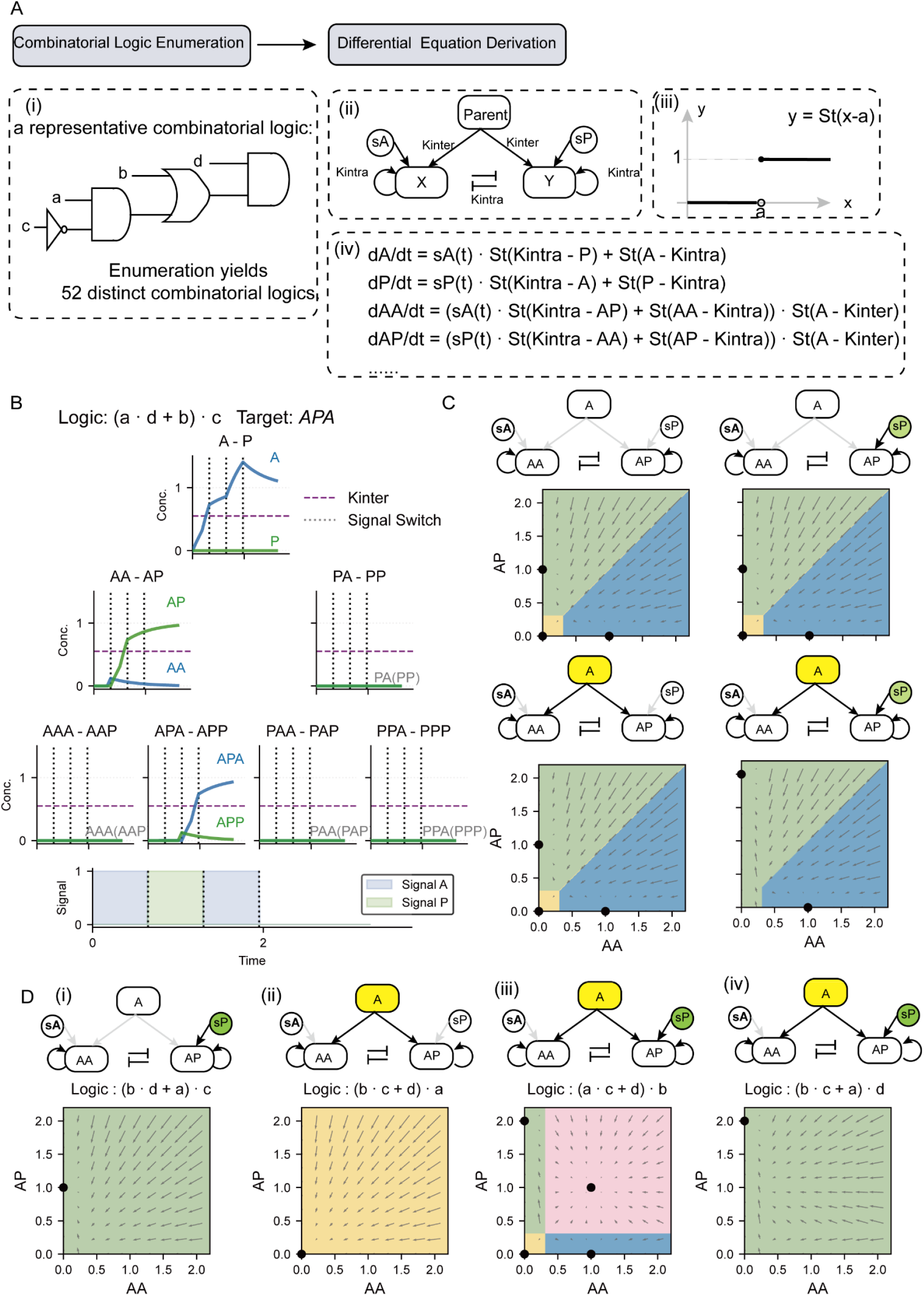
conserved attractor basin structure for targeted sequential fate induction. (A) Schematic of continuous differential equation derivation. (i): Equivalent logic gate circuit for a representative logic, and the enumeration yielded 52 distinct combinatorial logics. (ii): Schematic of CIS module with a parent node. *Kintra* is the threshold for self-activation and cross-inhibition (intra-module interaction), and *Kinter* is the threshold for parent activation (inter-module gating). (iii): Schematic of the Heaviside step function. (iv): The explicit ordinary differential equations derived from this representative logic shown in (i). (B) A successful dynamical simulation for targeted induction of *APA* fate. Blue and green curves represent transcription factors activated by signals sA and sP, respectively. Subplots in different rows represent transcription factors located at varying depths of the binary transcription factor tree. (C) Phase portraits (AA-AP) for a representative successful logic under different parent and signal conditions. In the accompanying network diagrams, gray arrows represent inactive parents or lack of signals, while colored nodes and black arrows indicate active parents or using inductive signals. Black spots denote stable attractors. Background colors represent different attractor basins: yellow for the dual-low progenitor state, blue and green for the A- and P-single-high differentiated states, and pink for dual-high state. This color scheme applies to all subsequent phase portraits. (D) Four typical aberrant phase portraits (AA-AP). (i) Phase portrait simulated with an inactive parent node and sP. (ii) Phase portrait simulated with an active parent node and no inductive signals. (iii) and (iv) Phase portraits simulated with an active parent node and sP.

To evaluate the 52 combinatorial logics, we performed a systematic parameter sweep by manually defining 10 sets of interaction thresholds (*Kintra* and *Kinter*) within the range of (0, 1). For each logic, we systematically scanned the signal switching period (Δ*t*_*s*_) to induce transcription factors at depth 6 of binary tree of interconnected CIS modules. The underlying ODEs were integrated to steady state, and the final expression levels were binarized to assess induction success based on the exact criteria from our initial Boolean screening. Ultimately, a combinatorial logic was deemed successful if proper induction was achieved for at least one Δ*t*_*s*_ under at least one (*Kintra, Kinter*) parameter pair (see Methods for details).

This screening revealed that six logic models achieved correct targeted induction under deterministic (noise-free) conditions, out of which only four remained strictly robust under additive stochastic noise perturbations (Figure S2). Notably, these four robust continuous models are identical to those identified in the Boolean enumeration. Their continuous trajectories faithfully mirror the Boolean cascade, exhibiting a strict sequential delay: downstream transcription factors remain functionally quiescent until the parent node concentration exceeds *Kinter*, at which point they become competent to respond to the inductive signal (Figure 3B).

Phase plane analysis reveals that all four successful logics share a conserved attractor basin structure, perfectly instantiating the discrete state-transition rules deduced earlier. Specifically, the system’s topology bifurcates around the inter-layer threshold (*Kinter*) (Figure 3C):

1. Parent-dependent gating: when the concentration of parent node is below *Kinter*, the uncommitted progenitor attractor persists regardless of the applied external signal.
2. Tri-stable landscape: when the concentration of parent node is above *Kinter*, and without inductive signals, the system needs to contain at least three attractors: an uncommitted progenitor attractor (low A, low P) and two differentiated attractors (high A or high P)
3. Signal-dependent fate selection and irreversibility: When the parent node is above *Kinter* and upon signal application, the system undergoes a directed bifurcation: the attractor basin for the targeted differentiated state expands to absorb the uncommitted progenitor basin, deterministically driving differentiation. Crucially, the basin of the alternative, non-targeted differentiated attractor remains stable, topologically ensuring commitment irreversibility.

Conversely, combinatorial logics violating these basin geometry constraints fail to execute targeted induction. The first class of failures violates parent-dependent gating rule: these logics destabilize the downstream progenitor cell attractor before the parent node concentration reaches *Kinter*, causing multiple layers to respond simultaneously rather than sequentially (Figure 3D(i)). The second mode of failure disrupts tri-stable landscape. Under non-inductive signal with an active parent node, these logics fail to maintain the necessary coexistence of the uncommitted progenitor attractor and the two differentiated attractors (Figure 3D(ii)). The third mode of failure involves aberrant attractor persistence or disappearance (Figure 3D(iii)-(iv)). Specifically, if the progenitor attractor fails to destabilize upon induction, the transcription factors remain trapped at low concentrations, rendering the cell unable to differentiate (Figure 3D(iii)). Conversely, if an established differentiated attractor erroneously disappears when exposed to opposing signals, the lineage commitment loses its topological irreversibility (Figure 3D(iv)).

We systematically categorized all failure modes of the 48 unsuccessful logics (Table S2, Figure S4), confirming that the conserved attractor basin structure identified above constitutes a necessary constraint for hierarchical induction.

### Precise signal switching period and non-inductive gap intervals jointly secure robust lineage induction at arbitrary depths

While the appropriate signal-induced landscape deformation satisfies the necessary requirement for hierarchical induction, it does not inherently guarantee correct temporal progression. If an external signal switches too rapidly, upstream transcription factors fail to reach the threshold required to unlock downstream layers. Conversely, if a signal persists for too long, the downstream layer prematurely perceives the input designated for the upstream tier, exhibiting unintended activation. For instance, under the signal sequence sA->sP (intended to sequentially activate TF A then TF AP), an overly prolonged sA pulse can erroneously activate the collateral TF AA. Successful hierarchical induction thus demands strict temporal coordination between external signal switching and intrinsic attractor landscape deformation. We therefore asked whether universal principles exist to guarantee precise induction.

We define the signal switching period Δ*t* as the duration of each inductive pulse (Figure 4A, middle panel) and investigate the existence of an optimal duration 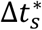 for induction at arbitrary lineage depths. Analytically, we prove that such an optimum universally exists provided the system satisfies the hierarchical threshold condition *Kintra* ≥ *Kinter* (Supplementary Note S2). Biologically, this condition dictates that a parent node must robustly consolidate its own lineage identity (governed by *Kintra*) before unlocking its downstream progeny (governed by *Kinter*). When an activated CIS module resting in the progenitor attractor gets induced by sA or sP, we define *t*_1_ as the time required for the activated TF to reach *Kintra*, and *t*_2_ the time to reach *Kinter* (Figure 4B). Under *Kintra* ≥ *Kinter*, the unique optimal switching period is exactly 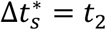. This implies that an idealized induction protocol should switch the signal the exact moment the active TF reaches the threshold to unlock the downstream layer. Both analytical derivations and numerical simulations confirm that 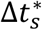 increases monotonically with *Kintra* and *Kinter* (Figure 4C; Figure S5), demonstrating that the optimal pulse duration is entirely determined by the system’s intrinsic kinetic parameters. This result provides a quantitative prescription for signal timing in hierarchical fate induction.

**Figure 4:**
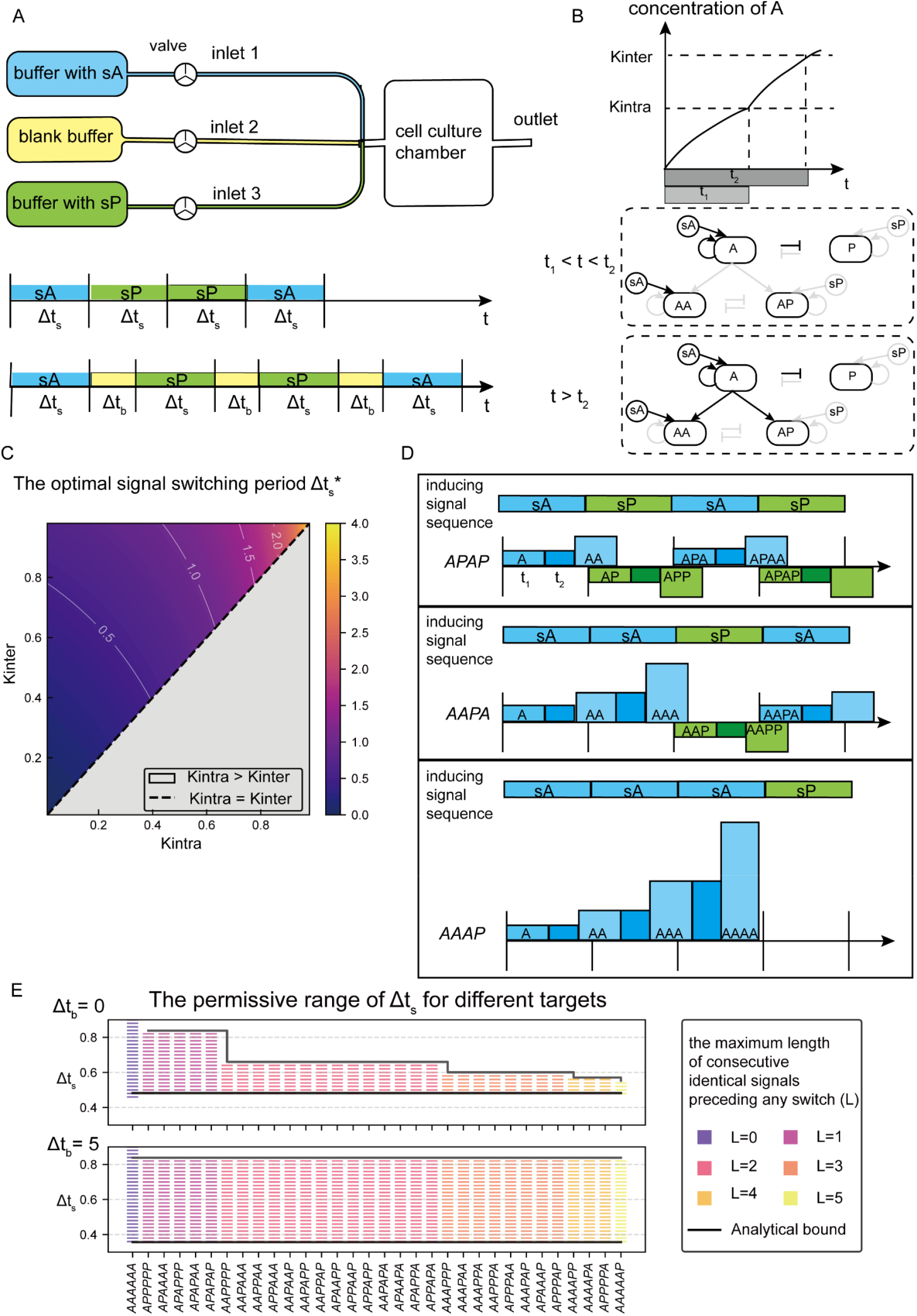
Analytical characterization of the signal switching period Δ*t*_*s*_ under the Heaviside step-function approximation. (A) Schematic of the induction protocol. Top: Using a microfluidic device to control the temporal sequence of external signals. Middle and Bottom: Definitions of the signal switching period Δ*t* and non-inductive gap interval Δ*t*_*s*_ are illustrated. (B) Schematic showing the definition and properties of *t*_1_ and *t*_2_. Light gray arrows represent inactive interactions while black arrows represent active interactions. Top: The light gray bar and dark gray bar indicates the definition of *t*_1_ and *t*_2_, respectively. Middle: When the concentration of A is above *Kintra* and below *Kinter*, A begins to self-activate and inhibit P. Bottom: When the concentration of A is above *Kinter*, A begins to activate its progenies. (C) Dependence of the theoretically optimal switching period 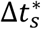 on *Kintra* and *Kinter*. The region where *Kintra* > *Kinter* (gray shading) admits no optimal switching period. In the region where *Kintra* < *Kinter*, the heatmap color indicates the magnitude of 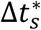. (D) Schematic of error accumulation induced by consecutive identical signals. Example target sequences of *L* = 1, 2, 3 (APAP, AAPA, and AAAP, respectively) are shown. Light blue and light green bars denote the time (*t*_1_) required for the target transcription factor (A- or P-terminal, respectively) in the current layer to accumulate from 0 to the intra-layer threshold Kintra. Dark blue and dark green bars represent the time (*t*_2_ − *t*_1_) required for these concentrations to increase from Kintra to Kinter. The vertical height of the rectangular bars indicates the increasing depth of layers when the signal remains unswitched. In the absence of signal switching, the temporal deviations (Δ*t*_*s*_ − *t*_2_) accumulate across layers. Consequently, in the AAAP sequence, the fourth-layer transcription factor erroneously responds to the third-layer signal, culminating in induction failure. (E) The permissive range of Δ*t*_*s*_ for different targets. Bar plots show the feasible Δ*t*_*s*_ ranges obtained from numerical simulation scans; different colors correspond to different values of the maximum length of consecutive identical signals preceding any switch (*L*). Red and black solid lines represent the analytically derived theoretical upper and lower bounds, respectively. The upper panel shows the permissive range or Δ*t*_*s*_ at Δ*t*_*b*_ = 0 ; the lower panel shows the case withΔ*t*_*b*_ = 5, where the permissive range of Δ*t*_*s*_ is significantly broadened.

While 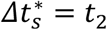defines the theoretical optimum, practical implementation requires understanding the permissive range of Δ*t*_*s*_. Further analysis reveals that the lower bound of this permissive range is exactly *t*_2_, indicating that an “switch immediately upon competence” strategy is always ideal. However, the upper bound is constrained by sequence-dependent leakage. Specifically, this kinetic limit is determined by the maximum length of consecutive identical signals preceding any switch, denoted as *L*. For example, the signal sequence sA->sA->sA->sP contains three identical pulses before the final switch, yielding *L* = 3. Sequences with larger *L* values exhibit a drastically narrowed permissive range of Δ*t*_*s*_. The mechanism of leakage accumulation can be conceptualized as a relay race: once an upstream TF reaches the inter-layer threshold at time *t*_2_, the downstream node begins to accumulate leaky expression for the remaining pulse duration, Δ*t*_*s*_−*t*_2_. During *L* consecutive identical signals, this advance leakage accumulates additively. To prevent the downstream node from falsely crossing its activation threshold (which intrinsically requires time *t*_1_), the total accumulated leakage time, *L*(Δ*t*_*s*_ − *t*_2_), must remain strictly below *t*_1_. This directly constrains the maximum allowable pulse duration to Δ*t*_*s*_ < *t*_2_ + *t*_1_/*L*. Fortunately, every alternating signal switch (e.g., sA to sP) acts as a dynamical reset, clearing out the accumulated leakage and preventing it from integrating further. Conversely, consecutive identical signals allow this leakage to integrate over time, eventually causing downstream nodes to prematurely cross activation thresholds and misinterpret the signal sequence (Figure 4D, Figure S6). This sequence-dependent narrowing dictates that deeper fates with homogeneous signaling histories are inherently more difficult to induce.

We next asked whether this sequence-dependent kinetic constraint could be systematically bypassed. Introducing a non-inductive gap interval (Δ*t*) between successive pulses provides a generalized structural solution (Figure 4A, bottom panel). Both analytical proofs and comprehensive numerical scanning demonstrate that increasing Δ*t*_*b*_ progressively broadens the permissive range of Δ*t*_*s*_, substantially lifting its upper limit (Figure S7). Intuitively, a sufficiently long signal withdrawal allows all residual, non-target leaky expression to naturally dissipate, effectively executing a memory reset before the next signal pulse. In this dissipative limit, the permissive range of Δ*t*_*s*_ expands from [*t*_2_, *t*_2_ + *t*_1_/*L*] to [*t*_1_, *t*_1_ + *t*_2_]. Consequently, the window width grows from *t*_1_/*L* to *t*_2_ and becomes completely independent of the sequence parameter *L* (Figure 4E; Figure S8). The introduction of Δ*t*_*b*_ thus transforms a fragile, sequence-dependent operating regime into a highly robust, sequence-independent one, ensuring that the induction protocol remains effective regardless of the target lineage.

### The perfect switching period requires idealized assumptions, but non-inductive gap intervals restore arbitrary-depth induction under realistic conditions

The optimal switching strategy derived above relies on two idealized assumptions: step-function regulation and symmetric parameters. However, real gene regulatory networks exhibit finite cooperativity, and the parameters governing X and Y in a CIS module are likely asymmetric, which naturally biases the binary fate decision[3, 28]. We therefore asked whether rules of optimal switching period 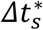 remain valid under these realistic conditions.

Our simulations show that under finite Hill cooperativity, an optimal switching period 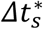 supporting arbitrary-depth induction no longer exists. Instead, the maximum inducible depth becomes finite (Figure 5A). This limitation stems from the inevitable leaky expression of downstream genes under continuous Hill functions (Figure 5B, Figure S8). In the previous idealized step-function model, a downstream module remains strictly locked at the (0,0) attractor as long as the upstream concentration stays below *Kinter*, ensuring a clean separation between inactive and responsive states. However, under Hill functions, the downstream attractor begins to deviate from (0,0) before the upstream concentration reaches *Kinter* (Figure 5C). In dynamical terms, finite cooperativity transforms strict gating into a continuous response. As a result, downstream transcription factors are prematurely activated. This baseline “leakiness” accumulates across successive layers, progressively threatening the fidelity of signal-directed induction and ultimately placing a ceiling on the induction depth.

**Figure 5:**
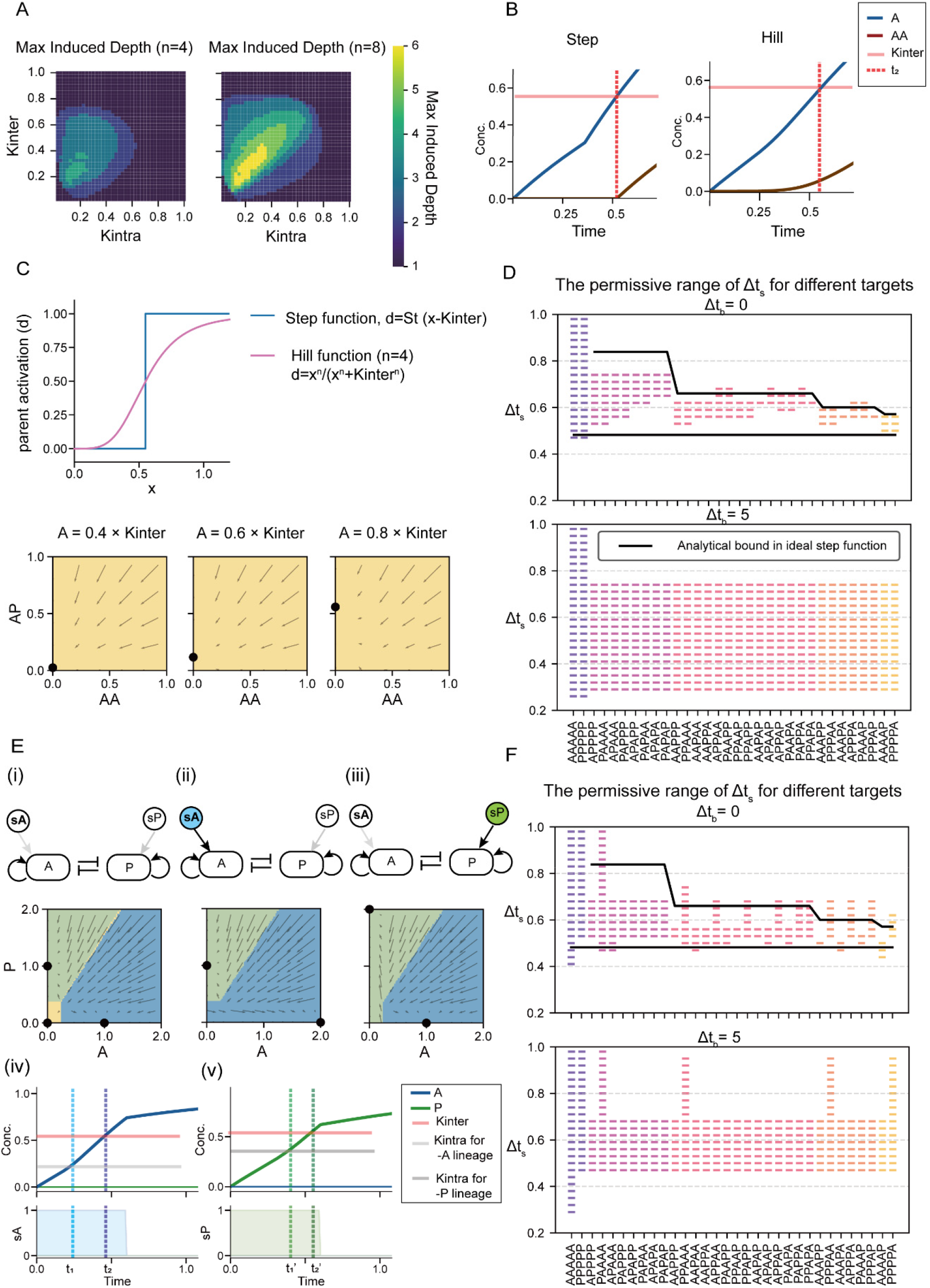
Induction of target cell fates under Hill kinetics and parameter asymmetry. (A) Maximum induction depth as a function of *Kintra* and *Kinter* in the absence of non-inductive gap intervals (Δ*t*_*b*_ = 0), shown for Hill coefficients n = 4, and 8 (left to right). Higher n values yield greater maximum depth, approaching the unlimited depth achievable under the step-function limit. (B) Dynamic simulations of targeted *AA* fate induction. Concentration kinetics of transcription factors A (blue curve) and AA (brown curve) are modeled using step (left) and Hill (right) functions. The horizontal pink line denotes *Kinter*. The vertical red dashed line marks *t*_2_, defined as the exact time point when the concentration of A reaches *Kinter*. Notably, under the step function model, AA expression initiates precisely at *t*_2_, while under Hill function, AA begins to express prior to A reaching Kinter. (C) Schematic and phase portraits demonstrating leaky expression under the Hill function. Top: Schematic of the Heaviside step function (blue line) and Hill function (pink line). The Horizontal axis denotes the concentration of parent node (x), and the vertical axis denotes the parent activation term (d). Under step function, *d* = *St* (*x* − *Kintra*) (defined as Figure 2A). Under Hill function, 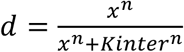. Bottom: Phase portraits of the AA-AP subsystem under Hill-function model (subject to signal sP). Parameters are set to Kintra=0.3 and Kinter=0.55. From left to right, the concentrations of the parent node (TF A) are 0.4×Kinter, 0.6×Kinter, and 0.8×Kinter, respectively (left to right). The black dots indicate the steady-state attractors, illustrating that under continuous Hill functions, the downstream attractor gradually deviates from zero even before the upstream concentration reaches Kinter. (D) Permissive range of Δ*t*_*s*_ for successful induction of each target sequence under Hill-function kinetics (n = 4). Bars show feasible Δ*t*_*s*_ from numerical scanning; colors indicate the maximum length of consecutive identical signals preceding any switch, *L*. black lines mark the analytical upper and lower bounds, derived from the step function model. Top panel: Δ*t*_*b*_ = 0 (without non-inductive gap intervals). Bottom panel: Δ*t*_*b*_ = 5, showing that non-inductive gap intervals substantially broaden the permissive windows. (E) Phase portraits and expression kinetics of the A–P subsystem under asymmetric parameters. (i-iii): Phase portraits of A-P subsystem. From left to right, the external signal conditions are blank (i), sA(ii), and sP(iii), respectively. Parameters: *Kintra*1 = 0.24 (governing Type-A auto-activation and A-to-P inhibition) and *Kintra*2 = 0.375 (governing Type-P auto-activation and P-to-A inhibition). The parameter asymmetry produces unequal attractor basins favoring the Type-A fate. (iv-v): Dynamic simulations of *A* fate and *P* fate induction over time. The blue and green solid curves represent the concentrations of transcription factors A and P, respectively. Horizontal lines denote the threshold: pink for Kinter, light gray for the *A*-lineage *Kintra*1 and dark gray for the *P*-lineage *Kintra*2. Vertical dashed lines mark the exact time points when the TF concentration reaches *Kintra* (*t*_1_ and *t*_1_′) and *Kinter* (*t*_2_ and *t*_2_′). Driven by the elevated Kintra threshold, Type-P fates require more time to escape the uncommitted progenitor basin (*t*_1_’ > *t*_1_) and activate downstream targets (*t*_2_’ > *t*_2_). (F) Permissive range of Δ*t*_*s*_ for successful induction of each target sequence under parameter asymmetry. Bars show feasible Δ*t* from numerical scanning; colors indicate the maximum length of consecutive identical signals preceding any switch, *L*. Top panel: Δ*t*_*b*_ = 0. Bottom panel: Δ*t*_*b*_ = 5. Comparison of the two panels shows that non-inductive gap intervals broaden the permissive window for all targets.

Despite this limitation, non-inductive gap intervals (Δ*t*_*b*_) restores robust induction across arbitrary depths (Figure 5D). Just as in the prior idealized case, this silent period broadens the permissive time window by allowing leaked downstream expression to decay back toward the (0,0) attractor. Given a sufficiently long Δ*t*_*b*_, the lower bound of Δ*t*_*s*_ decreases toward *t*_1_, the time required for parent-layer concentrations to exceed *Kintra* (though *t*_1_ can no longer be solved analytically under Hill kinetics). Therefore, while no single Δ*t*_*s*_ can achieve arbitrary-depth induction under Hill kinetics alone, introducing adequate non-inductive gap intervals fully recovers this capacity. A permissive Δ*t*_*s*_ range emerges where any target fate at any depth can be correctly induced.

A similar breakdown occurs when parameter symmetry is relaxed. When induction kinetics differ between competing transcription factors, the time required to escape the progenitor basin varies across lineages. For simplicity, we focus on a representative parameter asymmetry that biases induction, making Type-A fates (induced by sA) easier to induce than Type-P fates (induced by sP), reflecting unequally sized basins of attraction. Under these conditions, Type-P fates require more time to escape the (0,0) basin than Type-A fates (Figure 5E). Therefore, a single switching period Δ*t*_*s*_ cannot simultaneously meet the timing requirements for both pathways, and no optimal 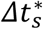 exists for unlimited hierarchical induction.

Once again, introducing non-inductive gap intervals (Δ*t*_*b*_) restores the network’s capacity for arbitrary-depth induction. By allowing sufficient relaxation between sequential pulses, Δ*t*_*b*_ compensates for timescale mismatches and prevents premature activation, allowing narrower but still achievable induction across all target sequences (Figure 5F, Note S3). Specifically, given a sufficiently long Δ*t*_*b*_, the allowable window for most target sequences converges to 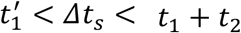. Here, the lower bound is set by the slower P-type kinetics 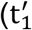 is the time for Type-P to reach its intra-layer threshold) and the upper bound is set by the faster A-type kinetics (*t*_1_, *t*_2_).

Together, these analyses reveal that the idealized perfect switching period 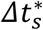 fails when either assumption (step-function regulation, parameter symmetry) is relaxed. However, the capacity for arbitrary-depth induction can be preserved in all cases when sufficiently long non-inductive gap intervals (Δ*t*_*b*_) are introduced, albeit with a narrowed permissive window.

### A polarization–division cycle provides a self-organized biological realization of temporal signal control to achieve hierarchical induction

Our prior analyses relied on externally imposed, precisely timed signal sequences. However, embryogenesis usually does not have centralized pacemakers or global external clocks[29, 30]. We therefore asked whether a multicellular system could autonomously generate the required spatiotemporal signal structure through local biophysical interactions,specifically both the inductive pulses and the reset intervals.

Inspired by the classic asymmetric cell divisions of early Caenorhabditis elegans embryogenesis, where *Wnt*/*β* − *catenin* signaling and anterior-posterior (A-P) polarity (via POP-1/SYS-1 segregation) sequentially specify lineage fates [31, 32], we constructed a 1D spatial model of self-organized polarization and division. The model architecture is governed by three local biophysical rules:

1. Intracellular polarity: Within each cell, two antagonistic membrane-associated proteins, a and p, self-recruit and mutually inhibit each other, spontaneously breaking symmetry to establish an internal A-P polarization axis.
2. Intercellular Ising-like coupling: Cells communicate exclusively via a short-range paracrine signal S (analogous to Wnt/MOM-2 relay). Signal S is secreted from the anterior pole (high a) and locally promotes the accumulation of protein p at the adjacent membrane of the neighboring cell. This interaction acts as a nearest-neighbor coupling term, aligning the polarization axes of adjacent cells in a head-to-tail fashion.
3. Orthogonal division and asymmetric inheritance: Cells divide along a cleavage plane orthogonal to their A-P polarization axis. Consequently, daughter cells inherit highly asymmetric quantities of proteins a and p. Following division, daughter cells enter a growth phase where they de novo synthesize proteins and dynamically re-establish membrane polarity before the next division event (Figure 6A). In this integrated framework, the inherited intracellular ratio of a versus p serves as the effective local sA or sP input to the downstream CIS gene regulatory network.

**Figure 6:**
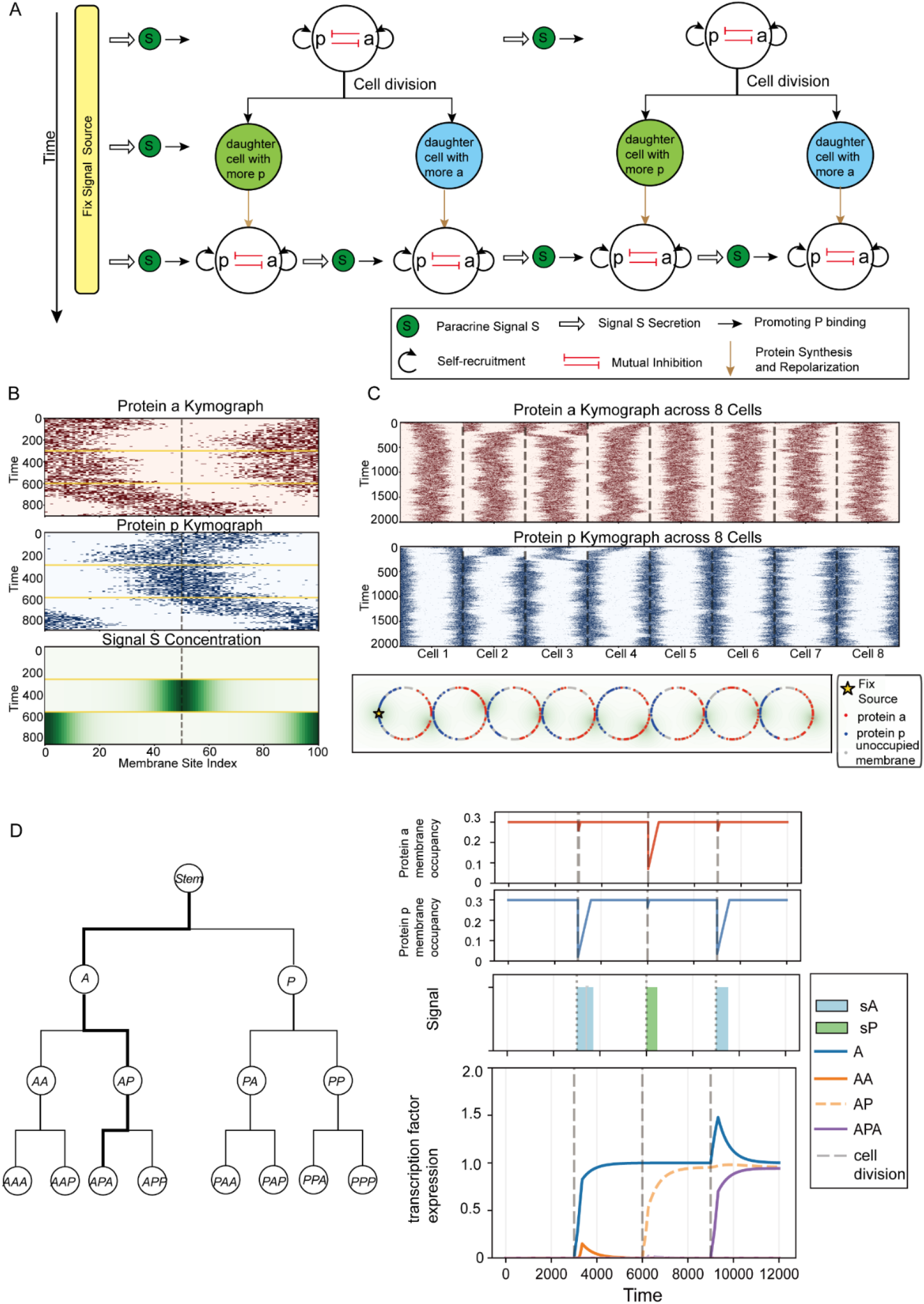
A self-organized polarization–division cycle biologically implements the temporal gating required for hierarchical induction. (A) Schematic of the polarization–division cycle. The black and red arrows represent the self-recruitment and mutual inhibition of proteins a and p, respectively, which drive spontaneous polarization. The paracrine signal S, denoted as deep green circles, is secreted from the anterior pole and aligns the polarity of adjacent cells. Upon division, asymmetric inheritance of a and p provides effective sA or sP inputs to the downstream transcription network. Blue and green circles represent daughter cells inherit more protein a and p, respectively. Following division, the protein synthesis and repolarization (brown arrows) act as a natural non-inductive gap interval (Δ*t*), resetting the system for the subsequent induction cycle. (B) Kymograph of polarity dynamics within a single cell. The cell first undergoes spontaneous symmetry breaking and polarization without external cues (*t* = 0 − 300 time units). Subsequent application of an external signal S to the right (*t* = 300 − 600) and then the left (*t* = 600 − 900) membrane demonstrates that the intracellular polarization axis can be dynamically reoriented by extracellular signals. (C) Kymograph showing long-range polarity alignment across a linear array of eight non-dividing cells. A fixed source of signal S at the left boundary mimics a localized embryonic organizing center. Local head-to-tail relay of signal S from each cell’s anterior pole is sufficient to propagate this directional cue, establishing a globally aligned anterior–posterior axis. (D) Lineage history and molecular dynamics of an *APA* fate cell. Left: Schematic of the lineage tree, with the bold black line tracing the history of a cell adopting the *APA* fate (Anterior -> Posterior -> Anterior daughters). Right: Corresponding simulated time-series dynamics for the tracked cell. Vertical dashed lines denote cell division events. The top two plots show the membrane occupancy of proteins a and p. The third plot shows the dynamics of effective signals (blue and green shaded regions for sA and sP, respectively). The fourth plot shows the hierarchical expression of the transcription factor network over time.

We first examined the spatial patterning of the system. Each individual cell, when not interacting with the neighbors, undergoes symmetry breaking spontaneously with polarity axes in random directions (Figure 6B). Once intercellular paracrine signaling S is active, local cell-to-cell communication acts as an effective ferromagnetic Ising coupling, rapidly propagating polarity orientation across the entire tissue chain, similar to prior works modeling the PCP cell patterning [33, 34]. Even without a global maternal gradient, short-range paracrine relay is sufficient to coordinate membrane polarity across long distances (Figure 6C). This confirms that local biophysical interactions can reliably construct a signaling scaffold for downstream fate specification.

We next incorporated cell division to evaluate the full spatiotemporal dynamics of the coupled system. Continuous division naturally drives the tissue through recurring cycles of polarization, asymmetric inheritance, and repolarization. When an aligned parent cell divides, the anterior daughter inherits a domain enriched in protein a, generating an immediate, effective pulse of signal sA to its internal transcriptional network; conversely, the posterior daughter receives an effective sP pulse. Crucially, immediately following division, the daughter cells are transiently unpolarized and must undergo a relaxation period to resynthesize proteins and re-establish their Ising alignment. During this physical repolarization phase, the inherited asymmetry degrades, and the effective intracellular inductive signals (sA and sP) decay below the inter-module gating threshold (*Kinter*). Of note, this relaxation period of the cell membrane acts as an intrinsic, system-generated non-inductive gap interval. Downstream transcription factor concentrations are temporarily relieved from external inductive drive, allowing kinetic leakage to safely dissipate toward the (0,0) attractor. Therefore, the biophysical time lag required for membrane repolarization directly coincides with, and physically implements, the dissipative non-inductive gap interval Δ*t_b_* required for robust hierarchical induction. As a result, the developing tissue autonomously unrolls the precise sequential, layer-by-layer induction cascade predicted by our theoretical framework, successfully navigating deep Waddington manifolds without requiring externally imposed pacemakers or global timing cues (Figure 6D).

This mechanism provides a concrete biophysical realization of our abstract control theory. While our initial analyses identified the stringent structural and kinetic constraints governing high-dimensional fate induction, this final model demonstrates how biology can elegantly satisfy these mathematical constraints during development. By cooperating the physical time scales of cell polarization and division, multicellular systems self-organize the precise “pulse-reset” signaling architecture required for robust, sequence-independent hierarchical fate specification.

## Discussion

In quantitative biology, the power of minimal modeling lies not in mirroring reality, but in identifying the failure modes and theoretical limits of biological mechanisms. By conceptualizing sequential fate specification as directed navigation across a Waddington manifold[2, 4, 35], we uncover a fundamental trade-off governing hierarchical induction under a minimalistic signaling alphabet: the conflict between commitment stability and inductive plasticity. To prevent new signals from overwriting established history, the attractor landscape must rigidly lock in upstream decisions; however, navigating toward deeper fates requires uncommitted downstream modules to remain kinetically plastic. We demonstrate that these two requirements create a severe operational bottleneck, especially under realistic kinetic constraints (such as continuous Hill regulation and parameter asymmetry)[3]. During consecutive pulses of identical signals, the plasticity required for downstream induction inevitably causes kinetic leakage in uncommitted layers. Meanwhile, the commitment stability transforms this premature leakage into irreversible off-target fate specification. Crucially, introducing non-inductive gap intervals (Δ*t*_*b*_) systematically decouples this trade-off. By allowing spurious leakage to dissipate without sacrificing upstream stability, non-inductive gap intervals restore accessibility and targeting precision across arbitrary lineage depths.

Recent advances in mathematical and data-driven modeling have made tremendous progress in quantifying Waddington’s epigenetic landscape, successfully mapping attractor topologies, non-equilibrium fluxes, and parameter bifurcations[35-37]. Yet, cell fate specification is not only about the existence of multiple attractors within a regulatory network, but also about the precise reachability of these states under biological constraints[38]. While these established frameworks provide a comprehensive picture of what cellular states exist and how parameters influence the steady-state landscape geometry, our study adds to these efforts by focusing on how a system actively navigates toward a pre-specified target fate when the external signaling repertoire is highly limited. Rather than conceptualizing differentiation as a passive relaxation toward a static equilibrium or as noise-driven diffusion between adjacent wells, our work models hierarchical fate specification as a problem of signal-directed transient dynamics: by continuously switching between a minimal set of discrete state-spaces under two or three external parameters (such as under a binary {sA, sP} or ternary {sA, sP, blank} input), the system uses temporal variation to guide the developmental trajectory, given appropriate coupling between its intrinsic timescale and the signaling timescale. From a non-equilibrium perspective, differentiation is thus steered by continuous transient kinetics rather than steady-state attraction alone.

Within this non-equilibrium framework, the non-inductive gap interval (Δ*t*_*b*_) acts as a critical kinetic reset, allowing “leakage” accumulation to dissipate before it triggers irreversible over-commitment. This theoretical necessity aligns with growing experimental evidence that developing cells actively decode temporal signaling dynamics[7]. For instance, it has been shown that the time-integral of BMP signaling dictates lineage bifurcation in embryonic stem cell models[39], while specific BMP pulse durations and sequences actively steer the binary choice between primitive streak and extraembryonic fates[40]. More important is the functional necessity of “silent” signaling intervals in some developmental processes. In mouse embryonic stem cells, Raina et al. observed that successive FGF/ERK pulses are separated by well-defined silent intervals that regulate downstream transcriptional processing[41]. Furthermore, recent optogenetic scanning of Wnt signaling identified an “anti-resonance” frequency filter, suggesting that the duration of inter-pulse OFF-times directly determines whether a tissue differentiation program is activated or if spurious, off-target activation is successfully filtered out[42]. Ultimately, these time-gating principles offer possible guidelines for synthetic biology: rather than relying on complex chemical cocktails, engineered circuits could achieve robust hierarchical differentiation by pairing hierarchical fate decision modules with timed pulse trains and mandatory washout intervals.

Beyond embryonic development, the strict requirement for coupling signaling timing with intrinsic cellular timescales may also offer dynamic insights into tissue repair and aging. Our model suggests that accurate lineage navigation relies on a precise temporal window of alternating inductive pulses (Δ*t*_*s*_) and non-inductive gap intervals (Δ*t*_*b*_). In early embryos, this well-timed oscillatory structure is naturally driven by active cell division and polarization cycles. However, in fully developed or aging tissues, the intrinsic biological hardware required to generate these periodic relaxation intervals is largely lost, as most specialized cells become post-mitotic or experience slowed cell-cycle dynamics. Consequently, if chronic environmental noise or injury pushes a cell into an aberrant phenotypic attractor (such as in pathological transdifferentiation or fibrosis), the absence of an appropriate signal sequence makes it difficult to redirect the cell back to its original identity. Moreover, without Δ*t*_*b*_ intervals to clear intracellular leakage, continuous therapeutic or corrective signaling may simply accumulate across multiple network layers, failing to overcome established commitment barriers and potentially exacerbating lineage mis-targeting. From this perspective, cellular aging may be linked to the intrinsic conflicts between stability and flexibility during embryogenesis.

Still, what is discussed in this work are highly simplified toy models. Our study relies on simplified combinatorial logics, deterministic dynamics, and an idealized one-dimensional cell chain model. In actual biological regulatory networks, multi-input integration is often more complex, and exogenous signals sometimes exert their effects by modulating intrinsic kinetic parameters (such as Hill activation thresholds) rather than acting as simple discrete logic inputs. Furthermore, while our supplementary stochastic simulations confirm the basic noise-robustness of this mechanism, real cellular environments exhibit strong intrinsic and extrinsic biochemical noise. Moreover, although our polarization-division model, inspired by membrane polarization and Wnt-mediated relay mechanisms in *C. elegans*, successfully demonstrates temporal signal reuse in a one-dimensional array, real embryogenesis involves much more intricate three-dimensional cellular architectures and morphogenetic dynamics. Beyond Wnt signaling, other contact-dependent pathways, such as Notch-Delta signaling, may also possess the biophysical capacity to drive similar polarization–division cycles. Therefore, incorporating multidimensional morphogenesis, strong stochasticity, and more heterogeneous regulatory functions into this framework constitutes a critical direction for future theoretical exploration.

In summary, this study provides a theoretical framework demonstrating how a limited set of signaling pathways, when temporally reused within specific GRN architectures, can generate a high diversity of cell fates. This finding not only offers a plausible theoretical resolution to the apparent paradox between signaling pathway minimalism and fate diversity observed in developmental processes such as those in *C. elegans*, but also provides key engineering design principles for synthetic biology. In engineered systems, CIS modules implementing appropriate combinatorial logic, together with temporally structured inductive and non-inductive signals, could enable targeted induction of arbitrary cell fates along a binary lineage tree. Furthermore, the construction of the polarization-division model suggests that, with appropriate upstream pathway design, such systems possess the potential to self-organize and autonomously acquire desired cell fates.

## Acknowledgement

This work was supported by the Fundamental and Interdisciplinary Disciplines Breakthrough Plan of the Ministry of Education of China (JYB2025XDXM502), National Key Research and Development Program of China (No. 2024YFA0919500), and National Natural Science Foundation of China (No. T2321001). LZ was supported in part by the Peking-Tsinghua Center for Life Sciences.

## Method

### Enumeration of Boolean regulatory logics

We represent the regulatory input received by each gene as a Boolean function of four elementary variables. For a given node *x*_*i*_, its state at discrete time *t* evolves according to

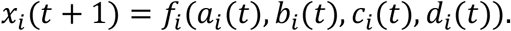

The four Boolean variables are defined as follows:

- *a*_*i*_(*t*): external signal matching, defined as *a*_*i*_(*t*) = 1 if the external signal *s*(*t*) matches node *i*, and 0 otherwise;
- *b*_*i*_(*t*) = *x*_*i*_(*t*): self-activation;
- *c*_*i*_(*t*) = 1 − *x*_sister(i)_(*t*): mutual inhibition from the sister node;
- *d*_*i*_(*t*) = *x*_parent(*i*)_(*t*): activation from the parent node. For first-layer nodes, *d*_*i*_(*t*) ≡ 1.

We enumerate all distinct regulatory logics as Boolean circuit topologies constructed from four inputs *a, b, c, d*. Each logic is represented as a rooted tree in which leaves correspond to input variables and internal nodes correspond to logical gates (AND or OR). Each input variable appears exactly once in the tree. To eliminate redundancies arising from circuit symmetries, we identify topologies up to isomorphism under the commutativity and associativity of logical operations. Using this representation, we exhaustively enumerate all non-equivalent circuit topologies, resulting in a total of 52 distinct regulatory logics.

### Boolean dynamical simulations and selection criteria

Boolean dynamics are simulated on a three-layer binary lineage tree. All nodes are initialized as inactive:

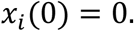

External signals are applied sequentially over four discrete time steps. Four target terminal states (AAA, AAP, APA, APP) are considered.

A candidate logic is classified as successful if it satisfies:

1. Target activation: the terminal node is active at the final time point;
2. Lineage integrity: all ancestor nodes are active;
3. Off-target suppression: all non-target nodes remain inactive.

### Mapping to continuous dynamical systems

To study the dynamical behavior beyond discrete updates, Boolean regulatory logics are mapped to continuous-time ordinary differential equations (ODEs). For each node *x*_*i*_, the dynamics are described by

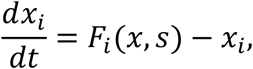

where *F*_*i*_(*x, s*) represents the combined regulatory input.

The external signal *s*_*i*_(*t*) is modeled as a binary, piecewise-constant input taking values in {0,1}, and is not transformed by a Hill function.

In contrast, endogenous regulatory interactions are modeled using Hill functions. Activation by a variable *y* is represented as

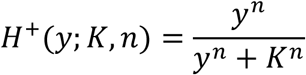

while inhibition is represented as

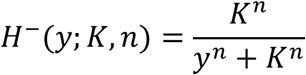

Accordingly, the regulatory inputs are mapped as:

- external signal: *a*_*i*_(*t*) → *s*_*i*_(*t*) ∈ {0,1},
- self-activation: *b*_*i*_(*t*) → *H*^+^(*x*_*i*_; *K*_intra_, *n*),
- mutual inhibition: *c*_*i*_(*t*) → *H*^-^(*x*_sister(i)_; *K*_intra_, *n*),
- upstream activation: *d*_*i*_(*t*) → *H*^+^(*x*_parent(i)_; *K*_inter_, *n*).

Boolean logical operations are mapped to continuous combinations as follows. Logical AND (∧) is implemented as multiplicative interaction:

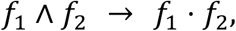

while logical OR (∨) is implemented as an additive combination:

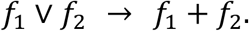

Using these rules, each Boolean logic *f*_*i*_(*a, b, c, d*) is translated into a continuous function *F*_*i*_(*x, s*) by replacing Boolean variables with their corresponding continuous representations.

As an illustrative example, the Boolean logic ((*a* ∧ *c*) ∨ *b*) ∧ *d* is mapped to

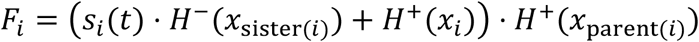

### Step-function approximation

We consider the limit *n* → ∞, under which Hill functions converge to step functions:

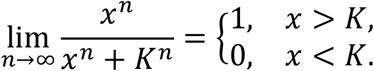

Adopting the step-function approximation and considering the combinatorial logic, the system reduces to a piecewise-linear switching system:

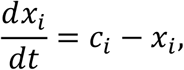

where *c*_*i*_ ∈ {0,1,2,3,4}.

### Induction protocol and success evaluation

To reduce computational cost, we restrict simulations to nodes along the target lineage and their direct antagonistic counterparts. This reduction does not affect the dynamics of the target trajectory, because the regulatory function *F*_*i*_(*x, s*) for each node depends only on its local inputs, namely its parent, its sister node, and itself. Nodes outside this local dependency set do not appear in the governing equations and therefore do not influence the evolution of the target lineage.

For each parameter set (*K*_intra_, *K*_inter_), we scan the signal switching period Δ*t*. For each Δ*t*, all distinct target sequences are simulated.

A trajectory is considered successful if the following conditions are satisfied at the end of induction:

i. the target node is activated relative to its sister node,

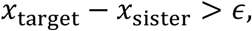
ii. all ancestor nodes along the target lineage are activated relative to their respective sister nodes,

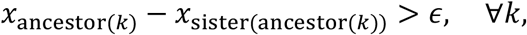

ensuring that the entire lineage path is consistently established.

Combinatorial logics and parameter sets achieving success rates above 80% are recorded.

### Basin of attraction analysis

The state space is discretized into a 200 × 200 grid. Each initial condition is numerically integrated until convergence.

Two trajectories are considered to reach the same attractor if their final states satisfy:

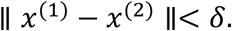

Vector fields are computed on a 20 × 20 grid for visualization.

### Symbolic analysis of expression dynamics

Under the step-function approximation, the system reduces to a piecewise-linear dynamical system. Within each segment, the governing equations take the form

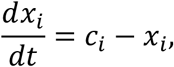

where *c*_*i*_ is constant within each segment.

The analytical solution is given by

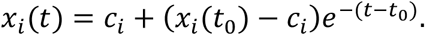

System evolution is partitioned by two types of events: (i) external signal switching at intervals Δ*t*; (ii) threshold-crossing events defined by *x*_*i*_(*t*) = *K*.

To construct full trajectories, we adopt a semi-symbolic, event-driven approach. At each segment, symbolic solutions are derived with symbolic initial conditions, while numerical evaluation is used to determine the ordering of competing events. Specifically, candidate event times are obtained by solving

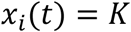

symbolically, and then evaluated numerically to identify the earliest event.

The system is then advanced to the next event time, and the final state of the current segment is used as the initial condition for the next segment. This procedure yields a globally consistent piecewise analytical trajectory.

This hybrid approach combines the accuracy of symbolic solutions with the robustness of numerical event detection, enabling exact characterization of switching times while avoiding ambiguities in event ordering.

### Parameter asymmetry

To investigate the effect of lineage bias, we introduce structured asymmetry into the regulatory thresholds. Rather than applying uniform perturbations, asymmetry is imposed in a direction-specific manner.

For A-biased configurations, threshold parameters associated with A-type regulation are modified as follows:

i. self-activation of A-type nodes is facilitated by decreasing the corresponding threshold,

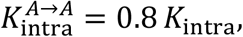
ii. inhibition of A-type nodes by their sister nodes is weakened by increasing the corresponding threshold,

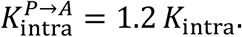

Conversely, for P-type nodes, the modifications are applied in the opposite direction:

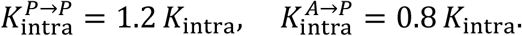

This parameterization introduces a systematic bias favoring A-type lineage commitment by simultaneously enhancing its self-activation and reducing its susceptibility to inhibition.

### Stochastic reaction–diffusion model of polarity establishment

We model the cell membrane as a one-dimensional periodic lattice representing a circular boundary of radius *R*, discretized into *L* evenly spaced sites. Periodic boundary conditions are imposed such that site indices are defined modulo *L*.

Each site *j* can be occupied independently by at most one molecule of protein *a* and one molecule of protein *p*, representing two antagonistic polarity species.

In addition to membrane-bound molecules, each cell maintains a cytoplasmic reservoir for both species. The total number of molecules is dynamically regulated, and the number of free molecules available for binding is given by the difference between the reservoir pool and membrane-bound molecules.

The system evolves through stochastic reaction–diffusion dynamics simulated using the Gillespie algorithm. At each lattice site, four classes of microscopic processes are considered for each protein species: membrane binding, unbinding, and lateral diffusion to adjacent sites.

Binding and unbinding rates depend not only on cytoplasmic availability but also on local membrane composition. Two nonlinear interaction mechanisms are incorporated:

- self-recruitment, whereby local enrichment enhances binding of the same species;
- mutual antagonism, whereby the opposing species suppresses binding and promotes unbinding.

Let *I*_*a*_(*j*) and *I*_*p*_(*j*) denote the local average occupancy of proteins *a* and *p* around site *j*, defined over a neighborhood of radius *ε*:

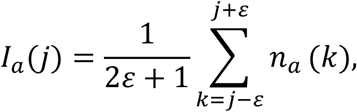

with periodic indexing, and similarly for *I*_*p*_(*j*). The propensity functions at site *j* are given by:

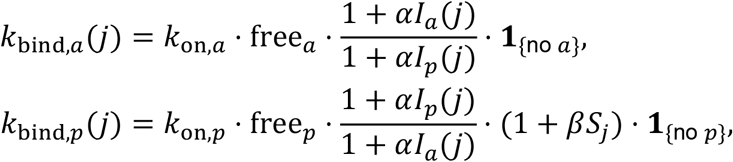

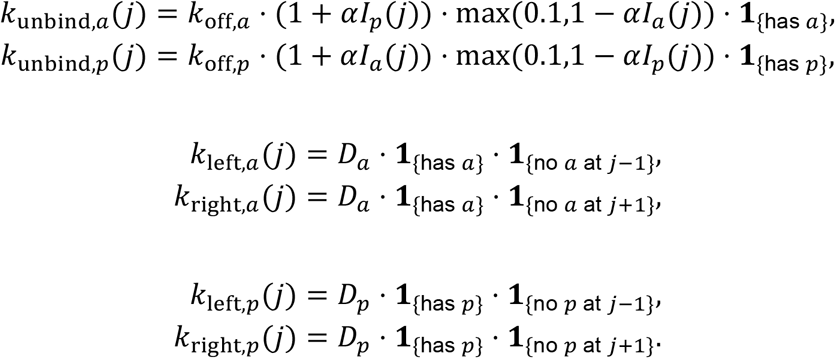

Here, *α* controls interaction strength, and *β* quantifies the modulation of protein *p* binding by the external signal field *S*_*j*_ (defined below).

At each step, all propensities are evaluated and reaction events are sampled using the Gillespie algorithm.

### Multicellular alignment

To model intercellular coupling, we consider a one-dimensional array of cells whose centers are arranged along a line with uniform spacing *d* ≥ 2*R*, where *R* is the cell radius. A fixed external signal source is placed at the origin.

Within each cell, polarity is defined by the spatial distribution of membrane-bound protein *a*. The polarity axis is computed as the average position of all membrane-bound *a* molecules, and the direction of this axis determines the location of signal emission. Upon polarization, each cell emits a signaling molecule *S* from its *a*-enriched pole.

We adopt a quasi-static approximation for signal propagation, in which the extracellular signal field is assumed to equilibrate instantaneously relative to intracellular dynamics. Under this assumption, we do not explicitly solve a diffusion equation; instead, each source contributes a steady-state concentration field given by

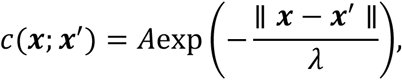

where ***x*** is the observation point, ***x***^′^ is the source location, and *λ* is the characteristic decay length.

The total signal concentration at position ***x*** is obtained by linear superposition of contributions from all sources:

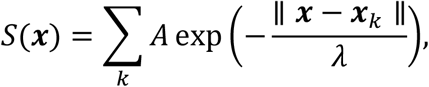

where ***x***_*k*_ denotes the emission point of the *K*-th source, including both the external source and polarized cells.

This formulation captures signal relay along the cell array, where polarized cells both respond to and reinforce the spatial signal gradient.

Signal sources are updated at discrete time intervals, while membrane dynamics evolve continuously via stochastic reactions. Between updates, the signal field is held constant. Each membrane site senses the local signal concentration based on its spatial position.

### Coupling polarity to gene expression

Cell division is incorporated as a discrete event occurring after a fixed duration *T*_div_. During division, membrane-bound proteins are partitioned according to their spatial positions, while cytoplasmic proteins are equally divided between the two daughter cells. The number of membrane binding sites *L* remains unchanged.

Protein production is introduced to maintain availability. When the total number of proteins (membrane-bound plus cytoplasmic) falls below a baseline level, new molecules are synthesized at a rate *K*_prod_.

Cellular polarity is quantified by the imbalance between the total amounts of proteins *a* and *p*. Based on this imbalance, the gene regulatory network receives a discrete input signal defined as:

- no signal if |*a*−*p*| < *θ*(*a* + *p*),
- *s*_*A*_ = 1 if *a* > *p* and the threshold is exceeded,
- *s*_*P*_ = 1 if *p* > *a* and the threshold is exceeded.

In all simulations, the threshold is set to *θ* = 0.2, representing the minimal asymmetry required to trigger fate specification.

## Notes

### Competing Interest Statement

The authors have declared no competing interest.

