## Supplementary material for "Directing hierarchical cell fate decisions through sequential pulses of minimal signaling alphabets": SI Appendix

Supplement

**Supplementary Tables**

**Table S1: Specific Expressions for the 52 Logic Gates**

| **Logic** | **Boolean Expression** |
| --- | --- |
| Logic 1 | $(a\times b+c)\times d$ |
| Logic 2 | $(a\times b+d)\times c$ |
| Logic 3 | $(a\times c+b)\times d$ |
| Logic 4 | $(a\times c+d)\times b$ |
| Logic 5 | $(a\times d+b)\times c$ |
| Logic 6 | $(a\times d+c)\times b$ |
| Logic 7 | $(a+b+c)\times d$ |
| Logic 8 | $(a+b+d)\times c$ |
| Logic 9 | $(a+b)\times(c+d)$ |
| Logic 10 | $(a+b)\times c\times d$ |
| Logic 11 | $(a+b)\times c+d$ |
| Logic 12 | $(a+b)\times d+c$ |
| Logic 13 | $(a+c+d)\times b$ |
| Logic 14 | $(a+c)\times(b+d)$ |
| Logic 15 | $(a+c)\times b\times d$ |
| Logic 16 | $(a+c)\times b+d$ |
| Logic 17 | $(a+c)\times d+b$ |
| Logic 18 | $(a+d)\times(b+c)$ |
| Logic 19 | $(a+d)\times b\times c$ |
| Logic 20 | $(a+d)\times b+c$ |
| Logic 21 | $(a+d)\times c+b$ |
| Logic 22 | $(b\times c+a)\times d$ |
| Logic 23 | $(b\times c+d)\times a$ |
| Logic 24 | $(b\times d+a)\times c$ |
| Logic 25 | $(b\times d+c)\times a$ |
| Logic 26 | $(b+c+d)\times a$ |
| Logic 27 | $(b+c)\times a\times d$ |
| Logic 28 | $(b+c)\times a+d$ |
| Logic 29 | $(b+c)\times d+a$ |
| Logic 30 | $(b+d)\times a\times c$ |
| Logic 31 | $(b+d)\times a+c$ |
| Logic 32 | $(b+d)\times c+a$ |
| Logic 33 | $(c\times d+a)\times b$ |
| Logic 34 | $(c\times d+b)\times a$ |
| Logic 35 | $(c+d)\times a\times b$ |
| Logic 36 | $(c+d)\times a+b$ |
| Logic 37 | $(c+d)\times b+a$ |
| Logic 38 | $a\times b\times c\times d$ |
| Logic 39 | $a\times b\times c+d$ |
| Logic 40 | $a\times b\times d+c$ |
| Logic 41 | $a\times b+c\times d$ |
| Logic 42 | $a\times b+c+d$ |
| Logic 43 | $a\times c\times d+b$ |
| Logic 44 | $a\times c+b\times d$ |
| Logic 45 | $a\times c+b+d$ |
| Logic 46 | $a\times d+b\times c$ |
| Logic 47 | $a\times d+b+c$ |
| Logic 48 | $a+b+c+d$ |
| Logic 49 | $b\times c\times d+a$ |
| Logic 50 | $b\times c+a+d$ |
| Logic 51 | $b\times d+a+c$ |
| Logic 52 | $c\times d+a+b$ |

**Table S2: Failure Reasons for the 52 Logics**

‘1’ indicates True, ‘0’ indicates False.

| **Logic** | **Saddle at** $S=(0,0)$ | **Only High-High Stem at** $S=(0,0)$ | **No Diff. State at**  $S=(0,0)$ | **Stem Persists under induction** | $<2$ **Diff. States under induction** | **Inter-layer Failure** |
| --- | --- | --- | --- | --- | --- | --- |
| 1 | 1 | 0 | 0 | 0 | 0 | 0 |
| 2 | 1 | 0 | 0 | 0 | 0 | 0 |
| 3 | 0 | 0 | 0 | 0 | 0 | 0 |
| 4 | 0 | 0 | 0 | 1 | 0 | 0 |
| 5 | 0 | 0 | 0 | 0 | 0 | 0 |
| 6 | 0 | 0 | 0 | 1 | 0 | 0 |
| 7 | 0 | 1 | 0 | 1 | 1 | 0 |
| 8 | 1 | 0 | 0 | 0 | 0 | 0 |
| 9 | 0 | 0 | 0 | 1 | 1 | 0 |
| 10 | 0 | 0 | 0 | 0 | 0 | 0 |
| 11 | 0 | 1 | 1 | 1 | 1 | 0 |
| 12 | 0 | 1 | 0 | 1 | 1 | 0 |
| 13 | 0 | 0 | 0 | 1 | 0 | 0 |
| 14 | 1 | 0 | 0 | 0 | 1 | 0 |
| 15 | 0 | 0 | 0 | 1 | 0 | 0 |
| 16 | 0 | 1 | 1 | 1 | 1 | 0 |
| 17 | 0 | 1 | 0 | 1 | 1 | 0 |
| 18 | 0 | 1 | 0 | 1 | 0 | 0 |
| 19 | 0 | 0 | 0 | 1 | 0 | 0 |
| 20 | 0 | 1 | 0 | 1 | 0 | 0 |
| 21 | 0 | 1 | 0 | 1 | 0 | 0 |
| 22 | 0 | 0 | 0 | 0 | 1 | 0 |
| 23 | 0 | 0 | 1 | 0 | 1 | 0 |
| 24 | 0 | 0 | 0 | 0 | 0 | 1 |
| 25 | 0 | 0 | 1 | 0 | 1 | 0 |
| 26 | 0 | 0 | 1 | 0 | 1 | 0 |
| 27 | 0 | 0 | 1 | 0 | 1 | 0 |
| 28 | 0 | 1 | 1 | 1 | 1 | 0 |
| 29 | 0 | 1 | 0 | 1 | 1 | 0 |
| 30 | 0 | 0 | 1 | 0 | 1 | 0 |
| 31 | 1 | 0 | 0 | 0 | 1 | 0 |
| 32 | 1 | 0 | 0 | 0 | 1 | 0 |
| 33 | 0 | 0 | 0 | 1 | 0 | 0 |
| 34 | 0 | 0 | 1 | 0 | 1 | 0 |
| 35 | 0 | 0 | 1 | 1 | 1 | 0 |
| 36 | 0 | 0 | 0 | 1 | 1 | 0 |
| 37 | 0 | 0 | 0 | 1 | 1 | 0 |
| 38 | 0 | 0 | 1 | 1 | 1 | 0 |
| 39 | 0 | 1 | 1 | 1 | 1 | 0 |
| 40 | 1 | 0 | 0 | 0 | 0 | 0 |
| 41 | 1 | 0 | 0 | 0 | 0 | 0 |
| 42 | 0 | 1 | 1 | 1 | 1 | 0 |
| 43 | 0 | 0 | 0 | 0 | 0 | 0 |
| 44 | 0 | 0 | 0 | 0 | 0 | 1 |
| 45 | 0 | 1 | 1 | 1 | 1 | 0 |
| 46 | 0 | 0 | 0 | 0 | 1 | 0 |
| 47 | 0 | 1 | 0 | 1 | 1 | 0 |
| 48 | 0 | 1 | 1 | 1 | 1 | 0 |
| 49 | 0 | 0 | 0 | 0 | 1 | 0 |
| 50 | 0 | 1 | 1 | 1 | 1 | 0 |
| 51 | 0 | 1 | 0 | 1 | 1 | 0 |
| 52 | 0 | 1 | 0 | 1 | 1 | 0 |

**Table S3: Gillespie Simulation Parameters**

| **Parameter** | **Physical Meaning** | **Value** |
| --- | --- | --- |
| $n_{1}$ | Total protein a is $n_{1}$ times the number of membrane binding sites | 0.3 |
| $n_{2}$ | Total protein p is $n_{2}$ times the number of membrane binding sites | 0.3 |
| $k_{on,a}$ | Binding rate constant of protein a to the membrane | 1 |
| $k_{on,p}$ | Binding rate constant of protein p to the membrane | 1 |
| $k_{off,a}$ | Dissociation rate constant of protein a from the membrane | 1.5 |
| $k_{off,p}$ | Dissociation rate constant of protein p from the membrane | 3 |
| $\alpha$ | Strength of self-recruitment and mutual inhibition | 5 |
| $\beta$ | Strength of signal-activated p binding | 5 |
| $D_{a}$ | Diffusion rate constant of protein a | 1 |
| $D_{p}$ | Diffusion rate constant of protein p | 1 |
| $\varepsilon$ | Range of self-recruitment and mutual inhibition | 10 |
| $L$ | Number of binding sites on the cell membrane | 100 |
| $R$ | Physical radius of the circular cell | 30 |
| $d$ | Distance between the centers of adjacent cells | 62 |
| $\lambda$ | Spatial decay characteristic length of the signal | 12 |
| $k_{production}$ | Synthesis rate of proteins a and p after division | 0.00025 |
| Duration | Time interval between two divisions | 3000 |
| $K_{intra}$ | Intra-layer Hill constant for transcription factors (A, P, AA, etc.) | 0.3 |
| $K_{inter}$ | Inter-layer Hill constant for transcription factors (A, P, AA, etc.) | 0.6 |
| $n$ | Hill coefficient for transcription factors (A, P, AA, etc.) | 6 |
| $k_{rate}$ | Expression rate of transcription factors | 0.001 |

**Supplementary notes**

**Note S1:** Boolean expressions and ordinary differential equations (ODEs) for A, P, AA, AP, PA, and PP of the four successful logics.

To study the dynamical behavior of the gene regulatory networks, we map the Boolean regulatory logics to continuous-time ordinary differential equations (ODEs). Let $[X]$ denote the continuous concentration of node $X$, where $X\in\{A,P,AA,AP,PA,PP\}$. The external signals are sA and sP. Nodes ending with ’A’ receive sA, while nodes ending with ’P’ receive sP.

Endogenous regulatory interactions are modeled using standard Hill functions with Hill coefficient $n$. We define Kintra as the half-activation/repression threshold for intra-layer interactions (self-activation and sister mutual inhibition), and Kinter as the half-activation threshold for inter-layer interactions (upstream parent activation). For first-layer nodes ($A$ and $P$), there is no upstream parent node, so the parent activation term is set to 1. The ODEs for the 6 transcription factors under the 4 successful logics are explicitly formulated as follows:

Logic 3: $(a\times c+b)\times d$

$$\begin{matrix} \frac{d\left[ A \right]}{dt} & =\left( \mathrm{sA}\frac{K_{\mathrm{intra}}^{n}}{K_{\mathrm{intra}}^{n}+[P]^{n}}+\frac{[A]^{n}}{K_{\mathrm{intra}}^{n}+[A]^{n}} \right)-\left[ A \right] & (S1.1) \\ \frac{d\left[ P \right]}{dt} & =\left( sP\frac{K_{\mathrm{intra}}^{n}}{K_{\mathrm{intra}}^{n}+[A]^{n}}+\frac{[P]^{n}}{K_{\mathrm{intra}}^{n}+[P]^{n}} \right)-\left[ P \right] & (S1.2) \\ \frac{d\left[ \mathrm{AA} \right]}{dt} & =\left( \mathrm{sA}\frac{K_{\mathrm{intra}}^{n}}{K_{\mathrm{intra}}^{n}+[AP]^{n}}+\frac{[AA]^{n}}{K_{\mathrm{intra}}^{n}+[AA]^{n}} \right)\frac{[A]^{n}}{K_{\mathrm{inter}}^{n}+[A]^{n}}-\left[ \mathrm{AA} \right] & (S1.3) \\ \frac{d\left[ \mathrm{AP} \right]}{dt} & =\left( \mathrm{sP}\frac{K_{\mathrm{intra}}^{n}}{K_{\mathrm{intra}}^{n}+[AA]^{n}}+\frac{[AP]^{n}}{K_{\mathrm{intra}}^{n}+[AP]^{n}} \right)\frac{[A]^{n}}{K_{\mathrm{inter}}^{n}+[A]^{n}}-\left[ \mathrm{AP} \right] & (S1.4) \\ \frac{d\left[ \mathrm{PA} \right]}{dt} & =\left( \mathrm{sA}\frac{K_{\mathrm{intra}}^{n}}{K_{\mathrm{intra}}^{n}+[PP]^{n}}+\frac{[PA]^{n}}{K_{\mathrm{intra}}^{n}+[PA]^{n}} \right)\frac{[P]^{n}}{K_{\mathrm{inter}}^{n}+[P]^{n}}-\left[ \mathrm{PA} \right] & (S1.5) \\ \frac{d\left[ \mathrm{PP} \right]}{dt} & =\left( \mathrm{sP}\frac{K_{\mathrm{intra}}^{n}}{K_{\mathrm{intra}}^{n}+[PA]^{n}}+\frac{[PP]^{n}}{K_{\mathrm{intra}}^{n}+[PP]^{n}} \right)\frac{[P]^{n}}{K_{\mathrm{inter}}^{n}+[P]^{n}}-\left[ \mathrm{PP} \right] & (S1.6) \end{matrix}$$

Logic 5: $(a\times d+b)\times c$

$$\begin{matrix} \frac{d\left[ A \right]}{dt} & =\left( sA+\frac{[A]^{n}}{K_{\mathrm{intra}}^{n}+[A]^{n}} \right)\frac{K_{\mathrm{intra}}^{n}}{K_{\mathrm{intra}}^{n}+[P]^{n}}-\left[ A \right] & (S1.7) \\ \frac{d\left[ P \right]}{dt} & =\left( sP+\frac{[P]^{n}}{K_{\mathrm{intra}}^{n}+[P]^{n}} \right)\frac{K_{\mathrm{intra}}^{n}}{K_{\mathrm{intra}}^{n}+[A]^{n}}-\left[ P \right] & (S1.8) \\ \frac{d\left[ \mathrm{AA} \right]}{dt} & =\left( \mathrm{sA}\frac{[A]^{n}}{K_{\mathrm{inter}}^{n}+[A]^{n}}+\frac{[AA]^{n}}{K_{\mathrm{intra}}^{n}+[AA]^{n}} \right)\frac{K_{\mathrm{intra}}^{n}}{K_{\mathrm{intra}}^{n}+[AP]^{n}}-\left[ \mathrm{AA} \right] & (S1.9) \\ \frac{d\left[ \mathrm{AP} \right]}{dt} & =\left( \mathrm{sP}\frac{[A]^{n}}{K_{\mathrm{inter}}^{n}+[A]^{n}}+\frac{[AP]^{n}}{K_{\mathrm{intra}}^{n}+[AP]^{n}} \right)\frac{K_{\mathrm{intra}}^{n}}{K_{\mathrm{intra}}^{n}+[AA]^{n}}-\left[ \mathrm{AP} \right] & (S1.10) \\ \frac{d\left[ \mathrm{PA} \right]}{dt} & =\left( \mathrm{sA}\frac{[P]^{n}}{K_{\mathrm{inter}}^{n}+[P]^{n}}+\frac{[PA]^{n}}{K_{\mathrm{intra}}^{n}+[PA]^{n}} \right)\frac{K_{\mathrm{intra}}^{n}}{K_{\mathrm{intra}}^{n}+[PP]^{n}}-\left[ \mathrm{PA} \right] & (S1.11) \\ \frac{d\left[ \mathrm{PP} \right]}{dt} & =\left( \mathrm{sP}\frac{[P]^{n}}{K_{\mathrm{inter}}^{n}+[P]^{n}}+\frac{[PP]^{n}}{K_{\mathrm{intra}}^{n}+[PP]^{n}} \right)\frac{K_{\mathrm{intra}}^{n}}{K_{\mathrm{intra}}^{n}+[PA]^{n}}-\left[ \mathrm{PP} \right] & (S1.12) \end{matrix}$$

Logic 10: $(a+b)\times c\times d$

$$\begin{matrix} \frac{d\left[ A \right]}{dt} & =\left( sA+\frac{[A]^{n}}{K_{\mathrm{intra}}^{n}+[A]^{n}} \right)\frac{K_{\mathrm{intra}}^{n}}{K_{\mathrm{intra}}^{n}+[P]^{n}}-\left[ A \right] & (S1.13) \\ \frac{d\left[ P \right]}{dt} & =\left( sP+\frac{[P]^{n}}{K_{\mathrm{intra}}^{n}+[P]^{n}} \right)\frac{K_{\mathrm{intra}}^{n}}{K_{\mathrm{intra}}^{n}+[A]^{n}}-\left[ P \right] & (S1.14) \\ \frac{d\left[ \mathrm{AA} \right]}{dt} & =\left( sA+\frac{[AA]^{n}}{K_{\mathrm{intra}}^{n}+[AA]^{n}} \right)\frac{K_{\mathrm{intra}}^{n}}{K_{\mathrm{intra}}^{n}+[AP]^{n}}\frac{[A]^{n}}{K_{\mathrm{inter}}^{n}+[A]^{n}}-\left[ \mathrm{AA} \right] & (S1.15) \\ \frac{d\left[ \mathrm{AP} \right]}{dt} & =\left( sP+\frac{[AP]^{n}}{K_{\mathrm{intra}}^{n}+[AP]^{n}} \right)\frac{K_{\mathrm{intra}}^{n}}{K_{\mathrm{intra}}^{n}+[AA]^{n}}\frac{[A]^{n}}{K_{\mathrm{inter}}^{n}+[A]^{n}}-\left[ \mathrm{AP} \right] & (S1.16) \\ \frac{d\left[ \mathrm{PA} \right]}{dt} & =\left( sA+\frac{[PA]^{n}}{K_{\mathrm{intra}}^{n}+[PA]^{n}} \right)\frac{K_{\mathrm{intra}}^{n}}{K_{\mathrm{intra}}^{n}+[PP]^{n}}\frac{[P]^{n}}{K_{\mathrm{inter}}^{n}+[P]^{n}}-\left[ \mathrm{PA} \right] & (S1.17) \\ \frac{d\left[ \mathrm{PP} \right]}{dt} & =\left( sP+\frac{[PP]^{n}}{K_{\mathrm{intra}}^{n}+[PP]^{n}} \right)\frac{K_{\mathrm{intra}}^{n}}{K_{\mathrm{intra}}^{n}+[PA]^{n}}\frac{[P]^{n}}{K_{\mathrm{inter}}^{n}+[P]^{n}}-\left[ \mathrm{PP} \right] & (S1.18) \end{matrix}$$

Logic 43: $a\times c\times d+b$

$$\begin{matrix} \frac{d\left[ A \right]}{dt} & =sA\frac{K_{\mathrm{intra}}^{n}}{K_{\mathrm{intra}}^{n}+[P]^{n}}+\frac{[A]^{n}}{K_{\mathrm{intra}}^{n}+[A]^{n}}-\left[ A \right] & (S1.19) \\ \frac{d\left[ P \right]}{dt} & =sP\frac{K_{\mathrm{intra}}^{n}}{K_{\mathrm{intra}}^{n}+[A]^{n}}+\frac{[P]^{n}}{K_{\mathrm{intra}}^{n}+[P]^{n}}-\left[ P \right] & (S1.20) \\ \frac{d\left[ \mathrm{AA} \right]}{dt} & =sA\frac{K_{\mathrm{intra}}^{n}}{K_{\mathrm{intra}}^{n}+[AP]^{n}}\frac{[A]^{n}}{K_{\mathrm{inter}}^{n}+[A]^{n}}+\frac{[AA]^{n}}{K_{\mathrm{intra}}^{n}+[AA]^{n}}-\left[ \mathrm{AA} \right] & (S1.21) \\ \frac{d\left[ \mathrm{AP} \right]}{dt} & =sP\frac{K_{\mathrm{intra}}^{n}}{K_{\mathrm{intra}}^{n}+[AA]^{n}}\frac{[A]^{n}}{K_{\mathrm{inter}}^{n}+[A]^{n}}+\frac{[AP]^{n}}{K_{\mathrm{intra}}^{n}+[AP]^{n}}-\left[ \mathrm{AP} \right] & (S1.22) \\ \frac{d\left[ \mathrm{PA} \right]}{dt} & =sA\frac{K_{\mathrm{intra}}^{n}}{K_{\mathrm{intra}}^{n}+[PP]^{n}}\frac{[P]^{n}}{K_{\mathrm{inter}}^{n}+[P]^{n}}+\frac{[PA]^{n}}{K_{\mathrm{intra}}^{n}+[PA]^{n}}-\left[ \mathrm{PA} \right] & (S1.23) \\ \frac{d\left[ \mathrm{PP} \right]}{dt} & =sP\frac{K_{\mathrm{intra}}^{n}}{K_{\mathrm{intra}}^{n}+[PA]^{n}}\frac{[P]^{n}}{K_{\mathrm{inter}}^{n}+[P]^{n}}+\frac{[PP]^{n}}{K_{\mathrm{intra}}^{n}+[PP]^{n}}-\left[ \mathrm{PP} \right] & (S1.24) \end{matrix}$$

**Note S2:** Derivation of analytical solutions.

Taking the analytical solution for expression dynamics of Logic 3, Target APA as an example.

Under the limit $n\to\infty$, the Hill functions in the governing equations reduce to Heaviside step functions $H(x)$. The system dynamics become piecewise-linear. To induce the target lineage **APA** (A $\to$ P $\to$ A), the external signals $(sA,sP)$ are applied in a sequence: $(1,0)$ for the first $\Delta$t, followed by $(0,1)$ for the next $\Delta$t, and finally $(1,0)$ to induce the terminal fate.

For the non-target nodes (i.e., $[P],[PA],[PP]$), their concentrations remain strictly zero throughout the entire process because their upstream activators never cross the required threshold $\mathrm{Kinter}$, or they are fully suppressed by the dominant sibling crossing Kintra.

The analytical solutions for the active nodes $([A],[AA],[AP],[APA],[APP])$ are partitioned into the following sequential event-driven phases:

Phase 1: Initial Induction of A ($0\leq t<t_{1}$)

- At $t=0$, signals are $S_{A}=1,S_{P}=0$ with all initial concentrations at zero.

$$\begin{matrix} \frac{d[A]}{dt} & =1-[A]\Longrightarrow[A](t)=1-e^{-t} & (S2.1) \end{matrix}$$

- This phase ends when $[A]$ crosses the intra-layer threshold $K_{\mathrm{intra}}$ at time $t_{1}$:

$$t_{1}=-\text{ln}\left( 1-K_{\mathrm{intra}} \right)$$

Phase 2: Self-Activation of A ($t_{1}\leq t<t_{2}$)

- Once $[A]>K_{\mathrm{intra}}$, the self-activation term turns on ($H([A]-K_{\mathrm{intra}})=1$).

$$\begin{matrix} \frac{d[A]}{dt} & =2-[A]\Longrightarrow[A](t)=2-(2-K_{\mathrm{intra}})e^{-(t-t_{1})} & (S2.2) \end{matrix}$$

- This phase ends when $[A]$ crosses the inter-layer threshold $K_{\mathrm{inter}}$ at time $t_{2}$:

$$t_{2}=t_{1}-\text{ln}\left( \frac{2-K_{\mathrm{inter}}}{2-K_{\mathrm{intra}}} \right)$$

Phase 3: Unlocking Layer 2 ($t_{2}\leq t<\Delta t$)

- For $t>t_{2}$, layer 2 is unlocked ($H([A]-K_{\mathrm{inter}})=1$). Since $S_{A}=1$, the off-target sibling $[AA]$ begins to accumulate.

$$\begin{matrix} [A](t) & =2-(2-K_{\mathrm{inter}})e^{-(t-t_{2})} & (S2.3) \\ \frac{d[AA]}{dt} & =1-[AA]\Longrightarrow[AA](t)=1-e^{-(t-t_{2})} & (S2.4) \end{matrix}$$

- To prevent $[AA]$ from auto-activating and locking the wrong fate, the first signal switch must occur before $[AA]$ crosses $K_{\mathrm{intra}}$. This yields the crucial constraint: $\Delta t-t_{2}<t_{1}\Longrightarrow\Delta t<t_{1}+t_{2}$. Let $A_{\Delta1}=[A](\Delta t)$ and $AA_{\Delta1}=[AA](\Delta t)$.

Phase 4: First Signal Switch and AP Induction ($\Delta t\leq t<t_{3}$)

- At $t=\Delta t$, signals switch to $S_{A}=0,S_{P}=1$. The node $[P]$ remains $0$ because $[A]>K_{\mathrm{intra}}$ suppresses its signal activation. $[AA]$ begins to decay, while the target $[AP]$ starts to grow.

$$\begin{matrix} [A](t) & =1-(1-A_{\Delta1})e^{-(t-\Delta t)} (K_{\mathrm{inter}}<1\text{, A remains }>K_{\mathrm{inter}}) & (S2.5) \\ [AA](t) & =AA_{\Delta1}e^{-(t-\Delta t)} & (S2.6) \\ \frac{d[AP]}{dt} & =1-[AP]\Longrightarrow[AP](t)=1-e^{-(t-\Delta t)} & (S2.7) \end{matrix}$$

- This phase ends when $[AP]$ crosses $K_{\mathrm{intra}}$ at $t_{3}$:

$$t_{3}=\Delta t-\text{ln}\left( 1-K_{\mathrm{intra}} \right)=\Delta t+t_{1}$$

Phase 5: Self-Activation of AP ($t_{3}\leq t<t_{4}$)

- $[AP]$ auto-activates, and $[AA]$ continues to decay to zero.

$$[AP]\begin{matrix} (t) & =2-(2-K_{\mathrm{intra}})e^{-(t-t_{3})} & (S2.8) \end{matrix}$$

- This phase ends when $[AP]$ crosses $K_{\mathrm{inter}}$ at $t_{4}$, unlocking Layer 3:

$$t_{4}=t_{3}-\text{ln}\left( \frac{2-K_{\mathrm{inter}}}{2-K_{\mathrm{intra}}} \right)$$

Phase 6: Unlocking Layer 3 ($t_{4}\leq t<2\Delta t$)

- Layer 3 is unlocked. Since $S_{P}=1$ is still active, the off-target sibling $[APP]$ starts to grow.

$$\begin{matrix} \left[ \mathrm{AP} \right]\left( t \right) & =2-\left( 2-K_{\mathrm{inter}} \right)e^{-\left( t-t_{4} \right)} & (S2.9) \\ \left[ \mathrm{APP} \right]\left( t \right) & =1-e^{-\left( t-t_{4} \right)} & (S2.10) \end{matrix}$$

Phase 7: Second Signal Switch and APA Induction ($2\Delta t\leq t<t_{5}$)

- At $t=2\Delta t$, signals switch back to $S_{A}=1,S_{P}=0$. $[APP]$ decays, and the terminal target $[APA]$ grows.

$$\begin{matrix} [APP](t) & =APP_{\Delta2}e^{-(t-2\Delta t)} & (S2.11) \\ [APA](t) & =1-e^{-(t-2\Delta t)} & (S2.12) \end{matrix}$$

- $[APA]$ crosses $K_{\mathrm{intra}}$ at $t_{5}=2\Delta t+t_{1}$.

Phase 8: Terminal Fate Lock ($t\geq t_{5}$)

$$[APA]\begin{matrix} (t) & =2-(2-K_{\mathrm{intra}})e^{-(t-t_{5})} & (S2.13) \end{matrix}$$

- $[APA]$ eventually crosses $K_{\mathrm{inter}}$ and asymptotically approaches a steady-state concentration of $2$, permanently locking the APA cell fate.

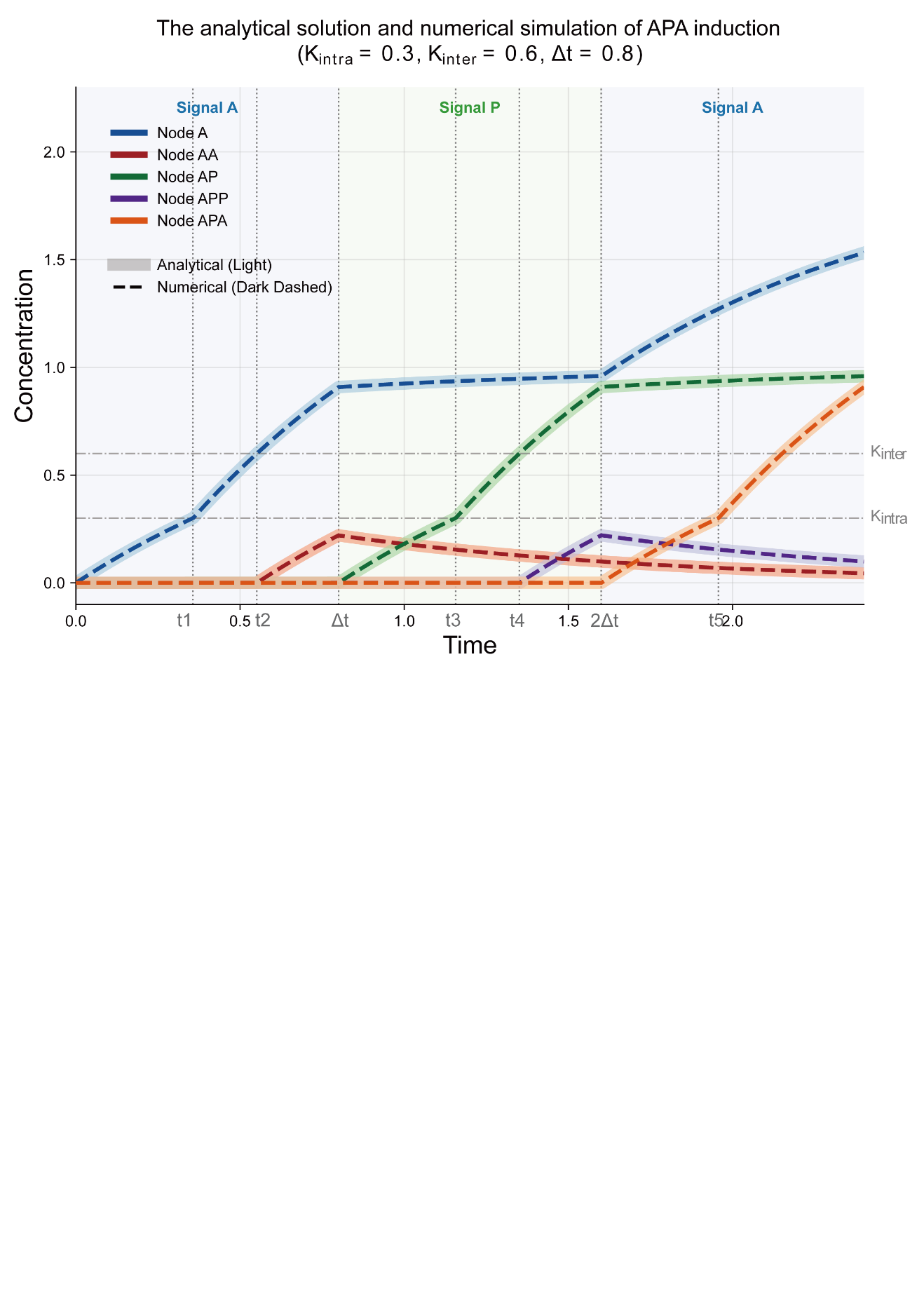

**Note S3:** Permissible $\Delta t$ under step functions with asymmetric parameters.

- To study the effect of asymmetric parameters, we mathematically formulate this A-biased configuration. The original uniform intra-layer threshold $K_{\mathrm{intra}}$ is bifurcated into node- and direction-specific parameters using a perturbation term $\Delta K>0$:
  - $K_{A,act}=K_{\mathrm{intra}}-\Delta K$: The self-activation threshold for A-type nodes. A decreased threshold facilitates their self-activation.
  - $K_{A,inh}=K_{\mathrm{intra}}+\Delta K$: The threshold for mutual inhibition exerted on A-type nodes by their sister nodes. An increased threshold weakens their susceptibility to inhibition.
  - $K_{P,act}=K_{\mathrm{intra}}+\Delta K$: The self-activation threshold for P-type nodes. An increased threshold hinders their self-activation.
  - $K_{P,inh}=K_{\mathrm{intra}}-\Delta K$: The threshold for mutual inhibition exerted on P-type nodes. A decreased threshold enhances their susceptibility to inhibition.
- Based on these definitions and taking Logic 3 as an example, the ordinary differential equations governing the A-biased asymmetric system are given by:

$$\begin{matrix} \frac{d[A]}{dt} & =\left( \mathrm{sA}\frac{K_{A,inh}^{n}}{K_{A,inh}^{n}+[P]^{n}}+\frac{[A]^{n}}{K_{A,act}^{n}+[A]^{n}} \right)-[A] & (S3.1) \\ \frac{d[P]}{dt} & =\left( \mathrm{sP}\frac{K_{P,inh}^{n}}{K_{P,inh}^{n}+[A]^{n}}+\frac{[P]^{n}}{K_{P,act}^{n}+[P]^{n}} \right)-[P] & (S3.2) \\ \frac{d[AA]}{dt} & =\left( \mathrm{sA}\frac{K_{A,inh}^{n}}{K_{A,inh}^{n}+[AP]^{n}}+\frac{[AA]^{n}}{K_{A,act}^{n}+[AA]^{n}} \right)\frac{[A]^{n}}{K_{\mathrm{inter}}^{n}+[A]^{n}}-[AA] & (S3.3) \\ \frac{d[AP]}{dt} & =\left( \mathrm{sP}\frac{K_{P,inh}^{n}}{K_{P,inh}^{n}+[AA]^{n}}+\frac{[AP]^{n}}{K_{P,act}^{n}+[AP]^{n}} \right)\frac{[A]^{n}}{K_{\mathrm{inter}}^{n}+[A]^{n}}-[AP] & (S3.4) \\ \frac{d[PA]}{dt} & =\left( \mathrm{sA}\frac{K_{A,inh}^{n}}{K_{A,inh}^{n}+[PP]^{n}}+\frac{[PA]^{n}}{K_{A,act}^{n}+[PA]^{n}} \right)\frac{[P]^{n}}{K_{\mathrm{inter}}^{n}+[P]^{n}}-[PA] & (S3.5) \\ \frac{d[PP]}{dt} & =\left( \mathrm{sP}\frac{K_{P,inh}^{n}}{K_{P,inh}^{n}+[PA]^{n}}+\frac{[PP]^{n}}{K_{P,act}^{n}+[PP]^{n}} \right)\frac{[P]^{n}}{K_{\mathrm{inter}}^{n}+[P]^{n}}-[PP] & (S3.6) \end{matrix}$$

- Under this asymmetric parameterization, the characteristic time $t_{1}$ and $t_{2}$ become distinct for A-type and P-type transcription factors, which we denote as $t_{1A}$, $t_{1P}$, $t_{2A}$, and $t_{2P}$ (where $t_{1A}<t_{1P}$ and $t_{2A}<t_{2P}$). Given a sufficiently long non-inductive interval $\Delta t_{2}$, the allowable window for the signal switching period $\Delta t$ can be analytically predicted depending on the target sequence:
  - For pure A-type lineages (all-A sequences), the allowable $\Delta t$ range is $[t_{1A},\infty)$.
  - For pure P-type lineages (all-P sequences), the allowable $\Delta t$ range is $[t_{1P},\infty)$.
  - For sequences involving exclusively P-to-A lineage switches (e.g., PAAAAA, PPAAAA, PPPAAA), the allowable $\Delta t$ range is $[t_{1P},t_{2P}+t_{1P})$.
  - For all other target sequences, the allowable $\Delta t$ range narrows to $[t_{1P},t_{2A}+t_{1A})$.
- The lower bound is uniformly dictated by $t_{1P}$, as the signal duration must be sufficient to ensure that even the slower P-type transcription factors can reach their $K_{\mathrm{intra}}$ threshold. The varying upper bounds arise from the premature accumulation of transcription factors in the non-induced layer. If a P-type transcription factor accumulates preemptively, it requires a time of $t_{2P}+t_{1P}$ to incorrectly cross its $K_{\mathrm{intra}}$. Conversely, because the A-type transcription factor accumulates faster, it only requires $t_{2A}+t_{1A}$ to reach its respective threshold, thereby imposing a stricter upper limit on the allowable signal duration.

**Note S4: Generalization of the System Dynamics Framework and High-Throughput Parameter Screening**

To further verify the universality of the discovered phase space topological constraints (specific attractor basin structures) in determining multi-stage cell fates, we generalized our dynamical modeling framework. In real biological systems, external signal regulation of target genes is rarely a simple Boolean switch (AND/OR gates); rather, it may operate by continuously altering biochemical parameters such as basal transcription rates or maximum reaction rates. Therefore, while retaining the core network motifs (the 8 combinatorial logics of auto-activation, mutual inhibition, and inter-layer activation), we allowed the external signal S to multi-dimensionally couple into the system equations via continuous weight parameters (c1, c2, c3, c4, c5).

Under this generalized framework, we performed Latin Hypercube Sampling (LHS) over the parameter space (v0, v1, v2, K, n, c1, c2, c3, c4, c5) to generate 2,000 parameter sets. Pairing these with the 8 topological logics, we constructed a total of 16,000 independent system models. Subsequently, we subjected these 16,000 systems to an extremely stringent topological screening of their static attractor basins. For the first layer (A-P phase diagram): Steady states were solved under S=0, 0.5, 1. The screening criteria dictated that at S=0.5, the system must contain at least one double-low stem cell attractor alongside A and P differentiation attractors; whereas upon applying S=0 or 1, the stem cell attractor must bifurcate and vanish, leaving only the two mutually exclusive differentiation attractors.

For the second layer (AA-AP phase diagram): To accurately capture the cascading effects, we computed 3 × 3 = 9 fine-grained phase diagrams. When the upstream node A is in the stem cell state, regardless of the value of S, the current stem cell attractor must persist. Conversely, when A is fully induced to its differentiation state (evaluated at its specific expression levels under S=0 and S=1, respectively), the phase diagram structure of the second layer must degenerate into a response pattern similar to that of the A-P phase diagram.

During the automated screening, classical Runge-Kutta integration is prone to stagnating at unstable saddle points on invariant manifolds, thereby generating "pseudo-attractors." To address this, we artificially introduced a single-step micro-perturbation during the late stages of simulated relaxation and prolonged the simulation time, which completely eliminated saddle-point misclassifications. Furthermore, to more authentically simulate the physiological starting point, we corrected the initial state for dynamic induction from the absolute zero origin to the genuine low-stem state, fully pre-equilibrated under the basal signal S=0.5.

Out of the 16,000 experiments, a total of 137 parameter sets perfectly passed the rigorous static basin screening described above. We then designated these 137 sets as the experimental group and randomly sampled 137 sets from the failed pool as the control group for continuous dynamic induction testing. We fixed the duration of blank gap interval at $\Delta t2=10$ and performed a fine-grained scan for the first signal duration $\Delta t$ within the interval (0, 100] (step size 0.1). If a parameter set possessed at least one viable $\Delta t$ window to successfully induce the cells along both the APAP and AAAP complex developmental trajectories to their final states, it was deemed a successful dynamic induction. The results demonstrated that within the 137 phase-diagram-screened experimental group sets, a remarkable 131 sets (95.6%) achieved successful dynamic induction; whereas within the 137 random control group sets, only 1 set (0.7%) exhibited the capacity for successful induction. This highly significant statistical divergence conclusively proves that: the specific basin morphology of the static phase space is not merely a necessary condition for achieving complex lineage development, but rather the strongest predictive hallmark for the success of multi-stage dynamic induction.

To explicitly illustrate the generalized framework discussed above, the complete mathematical formulations for the eight combinatorial logic gates are detailed as follows.

We first define the universal modular functions representing basal expression ($V_{\text{basal}}$), self-activation ($F_{\text{act}}$), mutual inhibition ($F_{\text{inh}}$), and inter-layer activation ($F_{\text{inter}}$).

$$\begin{matrix} V_{\text{basal}}(S) & =v_{0}\left( 1-c_{5}/2+c_{5}S \right) & (S4.1) \\ F_{\text{act}}(X,S) & =\frac{v_{1}(1-c_{1}/2+c_{1}S)X^{n}}{\frac{K^{n}}{(1-c_{2}/2+c_{2}S)}+X^{n}} & (S4.2) \\ F_{\text{inh}}(Y,S) & =\frac{v_{2}(1-c_{3}/2+c_{3}S)(1-c_{4}/2+c_{4}S)K^{n}}{Y^{n}+K^{n}(1-c_{4}/2+c_{4}S)} & (S4.3) \\ F_{\text{inter}}(Z) & =\frac{Z^{n}}{Z^{n}+K_{\text{inter}}^{n}} & (S4.4) \end{matrix}$$

**Layer 1: A and P Equations**

Because Layer 1 nodes do not receive upstream signals (i.e., component $d$ is absent), the 8 logic gates collapse into two shared topological cores based on how $b$ and $c$ interact.

**Core 1: OR-Gate Integration (b+c) (Shared by Logics 1, 2, 3, and 4)**

$$\begin{matrix} \frac{dA}{dt} & =V_{\text{basal}}\left( S \right)+F_{\text{act}}\left( A,S \right)+F_{\text{inh}}\left( P,S \right)-A & (S4.5) \\ \frac{dP}{dt} & =V_{\text{basal}}\left( 1-S \right)+F_{\text{act}}\left( P,1-S \right)+F_{\text{inh}}\left( A,1-S \right)-P & (S4.6) \end{matrix}$$

**Core 2: AND-Gate Integration (b×c)(Shared by Logics 5, 6, 7, and 8)**

$$\begin{matrix} \frac{dA}{dt} & =V_{\text{basal}}\left( S \right)+F_{\text{act}}\left( A,S \right)\times F_{\text{inh}}\left( P,S \right)-A & (S4.7) \\ \frac{dP}{dt} & =V_{\text{basal}}\left( 1-S \right)+F_{\text{act}}\left( P,1-S \right)\times F_{\text{inh}}\left( A,1-S \right)-P & (S4.8) \end{matrix}$$

**Layer 2: AA and AP Equations**

For Layer 2, the nodes $X\in\{AA,AP\}$ receive regulatory input from their upstream parent $Z=A$. The 8 possible canonical topologies combining $b$, $c$, and $d$ are expanded below.

**Group 1: Derived from OR-Gate Core (**$\boldsymbol{b}\mathbf{+}\boldsymbol{c}$**)**

**Logic 1:** $d\times(b+c)$

$$\begin{matrix} \frac{d[AA]}{dt} & =V_{\text{basal}}(S)+\left[ F_{\text{act}}([AA],S)+F_{\text{inh}}([AP],S) \right]\times F_{\text{inter}}(A)-[AA] & (S4.9) \\ \frac{d[AP]}{dt} & =V_{\text{basal}}(1-S)+\left[ F_{\text{act}}([AP],1-S)+F_{\text{inh}}([AA],1-S) \right]\times F_{\text{inter}}(A)-[AP] & (S4.10) \end{matrix}$$

**Logic 2:** $b\times d+c$

$$\begin{matrix} \frac{d[AA]}{dt} & =V_{\text{basal}}(S)+F_{\text{act}}([AA],S)\times F_{\text{inter}}(A)+F_{\text{inh}}([AP],S)-[AA] & (S4.11) \\ \frac{d[AP]}{dt} & =V_{\text{basal}}(1-S)+F_{\text{act}}([AP],1-S)\times F_{\text{inter}}(A)+F_{\text{inh}}([AA],1-S)-[AP] & (S4.12) \end{matrix}$$

**Logic 3:** $c\times d+b$

$$\begin{matrix} \frac{d[AA]}{dt} & =V_{\text{basal}}(S)+F_{\text{inh}}([AP],S)\times F_{\text{inter}}(A)+F_{\text{act}}([AA],S)-[AA] & (S4.13) \\ \frac{d[AP]}{dt} & =V_{\text{basal}}(1-S)+F_{\text{inh}}([AA],1-S)\times F_{\text{inter}}(A)+F_{\text{act}}([AP],1-S)-[AP] & (S4.14) \end{matrix}$$

**Logic 4:** $b+c+d$

$$\begin{matrix} \frac{d[AA]}{dt} & =V_{\text{basal}}(S)+F_{\text{act}}([AA],S)+F_{\text{inh}}([AP],S)+F_{\text{inter}}(A)-[AA] & (S4.15) \\ \frac{d[AP]}{dt} & =V_{\text{basal}}(1-S)+F_{\text{act}}([AP],1-S)+F_{\text{inh}}([AA],1-S)+F_{\text{inter}}(A)-[AP] & (S4.16) \end{matrix}$$

**Group 2: Derived from AND-Gate Core (b×c)**

**Logic 5:** $b\times c\times d$

$$\begin{matrix} \frac{d[AA]}{dt} & =V_{\text{basal}}(S)+F_{\text{act}}([AA],S)\times F_{\text{inh}}([AP],S)\times F_{\text{inter}}(A)-[AA] & (S4.17) \\ \frac{d[AP]}{dt} & =V_{\text{basal}}(1-S)+F_{\text{act}}([AP],1-S)\times F_{\text{inh}}([AA],1-S)\times F_{\text{inter}}(A)-[AP] & (S4.18) \end{matrix}$$

**Logic 6:** $(b+d)\times c$

$$\begin{matrix} \frac{d[AA]}{dt} & =V_{\text{basal}}(S)+\left[ F_{\text{act}}([AA],S)+F_{\text{inter}}(A) \right]\times F_{\text{inh}}([AP],S)-[AA] & (S4.19) \\ \frac{d[AP]}{dt} & =V_{\text{basal}}(1-S)+\left[ F_{\text{act}}([AP],1-S)+F_{\text{inter}}(A) \right]\times F_{\text{inh}}([AA],1-S)-[AP] & (S4.20) \end{matrix}$$

**Logic 7:** $(c+d)\times b$

$$\begin{matrix} \frac{d[AA]}{dt} & =V_{\text{basal}}(S)+\left[ F_{\text{inh}}([AP],S)+F_{\text{inter}}(A) \right]\times F_{\text{act}}([AA],S)-[AA] & (S4.21) \\ \frac{d[AP]}{dt} & =V_{\text{basal}}(1-S)+\left[ F_{\text{inh}}([AA],1-S)+F_{\text{inter}}(A) \right]\times F_{\text{act}}([AP],1-S)-[AP] & (S4.22) \end{matrix}$$

**Logic 8:** $b\times c+d$

$$\begin{matrix} \frac{d[AA]}{dt} & =V_{\text{basal}}(S)+F_{\text{act}}([AA],S)\times F_{\text{inh}}([AP],S)+F_{\text{inter}}(A)-[AA] & (S4.23) \\ \frac{d[AP]}{dt} & =V_{\text{basal}}(1-S)+F_{\text{act}}([AP],1-S)\times F_{\text{inh}}([AA],1-S)+F_{\text{inter}}(A)-[AP] & (S4.24) \end{matrix}$$

**Supplementary Figures**

**Figure S1:** State transition diagrams of the four successful logics.

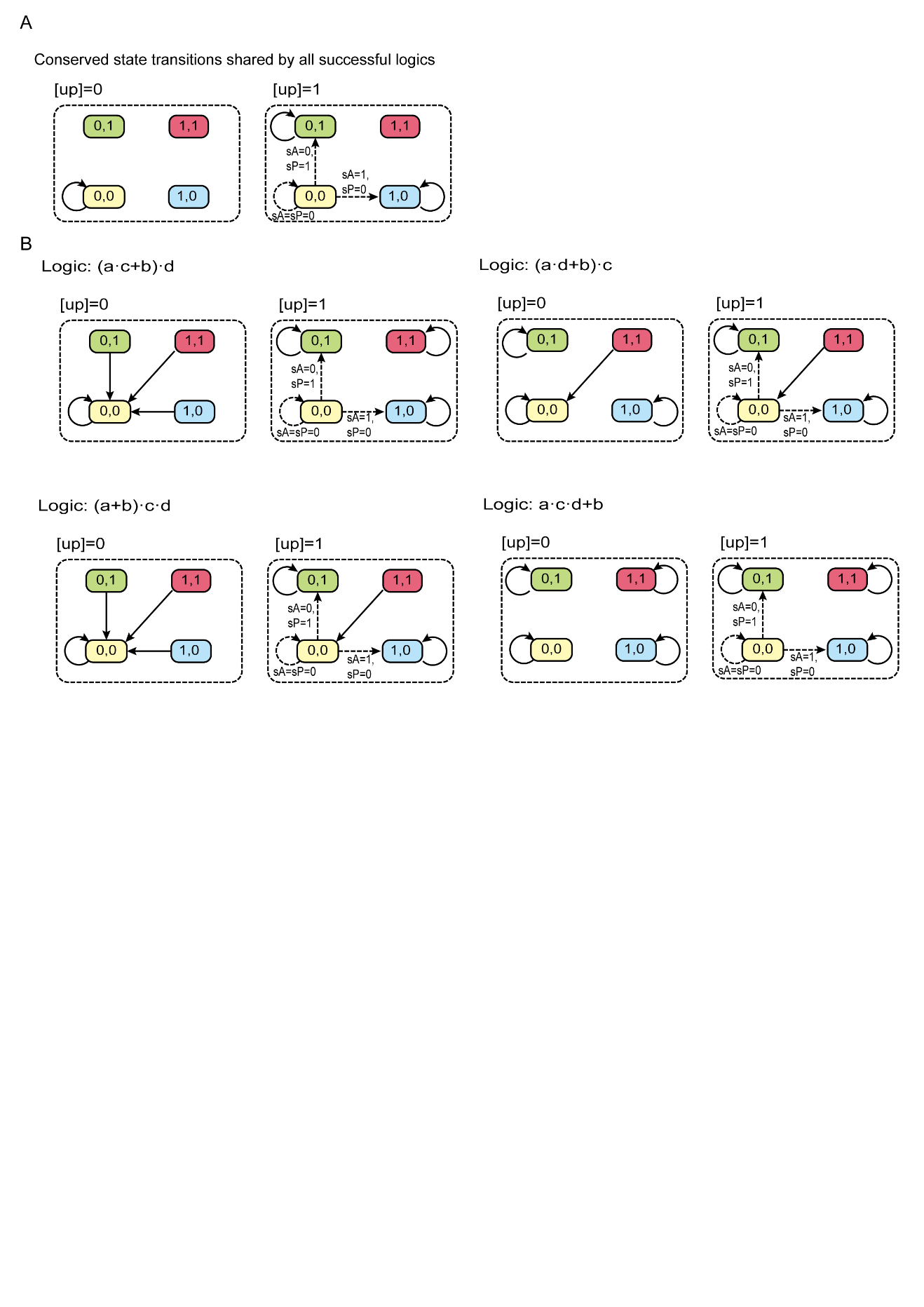

(A) Conserved state transition edges shared by all four successful Boolean logics. Square nodes represent the four distinct cellular states: double-low stem cell state (yellow), xP-high differentiated state (green), xA-high differentiated state (blue), and double-high mixed state (red). Solid arrows denote state transitions that are unconditionally present, whereas dashed arrows denote transitions that occur only under specific external signal inputs (e.g., sA=0 and sP=1).

(B) Distinct state transition topologies unique to each of the four successful logics.

**Figure S2:** Dynamics simulations of two noise-susceptible logics and a robust control.

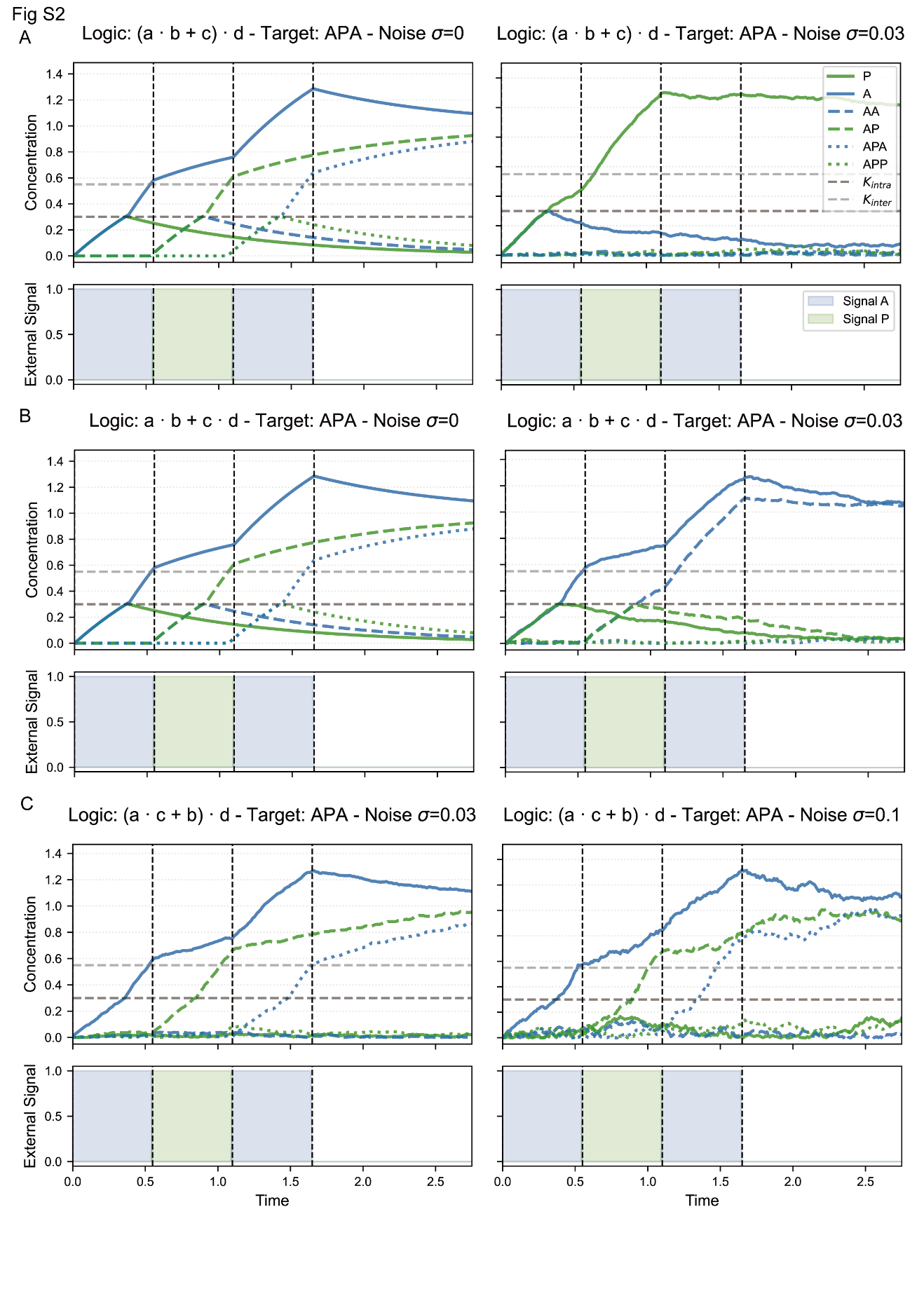

(A-B) Dynamic simulations of two logic configurations that successfully induce the target lineage (APA) in the absence of noise ($\sigma=0$, left panels), but are highly susceptible to little noise perturbations. Blue and green curves represent transcription factors responsive to external signals sA and sP, respectively. Solid, dashed, and dotted lines denote factors in the first, second, and third layers of the regulatory network, respectively. As shown in the right panels, even a slight noise level ($\sigma=0.03$) is sufficient to disrupt precise temporal induction by triggering the ectopic activation of off-target lineages (e.g., the rapid rise of green curves).

(C) Dynamic simulations of a noise-robust logic. This logic configuration maintains accurate temporal induction of the target lineage (APA) without off-target leakage, under low ($\sigma=0.03,$left panel) and moderate ($\sigma=0.1$, right panel) noise conditions.

**Figure S3:** A-P, AA-AP phase diagrams of the four successful logics. (Related to Figures 2C and 2D).

While Figure 2 illustrates the phase space topology for a representative logic configuration, this figure comprehensively presents the complete set of A-P (first layer) and AA-AP (second layer) phase diagrams for all four successful logic combinations. Two-dimensional phase portraits and attractor basins are shown under varying external signal combinations (sA, sP) and upstream node concentrations (e.g., A=0.54 or A=0.56). Consistent with Figure 2, background colors denote distinct basins of attraction representing specific cell fates: double-low stem cell state (yellow), xA-high differentiated state (blue), xP-high differentiated state (green), and double-high mixed state (red). White arrows indicate the vector fields, and black open circles mark the locations of attractors.

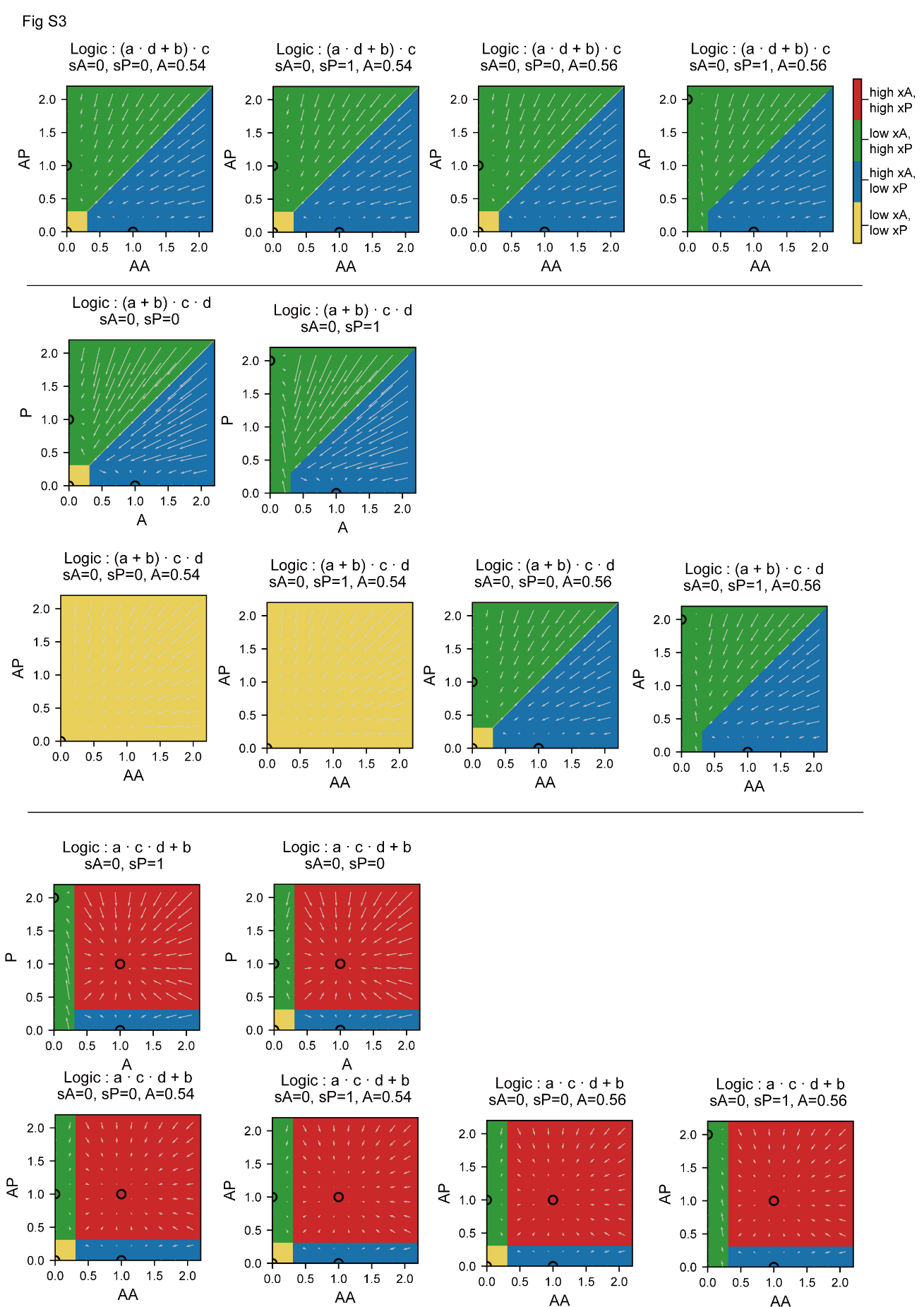

**Figure S4:** Detailed demonstration of failed logics, including failure phase diagrams and failed dynamics simulations. (Related to Figures 2E and 2F).

This figure provides a comprehensive catalog of six representative failed logic configurations. For each logic, the static basins of attraction (top) and the corresponding temporal dynamics during an attempted induction of the APA lineage (bottom) are evaluated.

Phase Diagrams (Top): For each logic, two A-P phase diagrams (layer 1) and four AA-AP phase diagrams (layer 2) are shown under various external signal and upstream node conditions. Consistent with previous figures, background colors denote the distinct basins of attraction (yellow: double-low stem state; blue: xA-high state; green: xP-high state; red: double-high mixed state). These failure cases typically exhibit topological defects in their phase space.

Dynamic Simulations (Bottom): Concentration trajectories are arranged in a hierarchical tree layout for clarity. Instead of using different line styles to distinguish network layers, the hierarchical levels are naturally represented by the spatial arrangement of the subpanels: the first (A vs P), second (AA vs AP, PA vs PP), and third layers are positioned in the top, middle, and bottom rows of the grid, respectively. Blue and green solid curves denote the A-terminal and P-terminal transcription factors. The external signal sequence (sA followed by sP, then sA) is displayed at the bottom. The horizontal dashed lines mark the intra-layer (Kintra) and inter-layer (Kinter) thresholds.

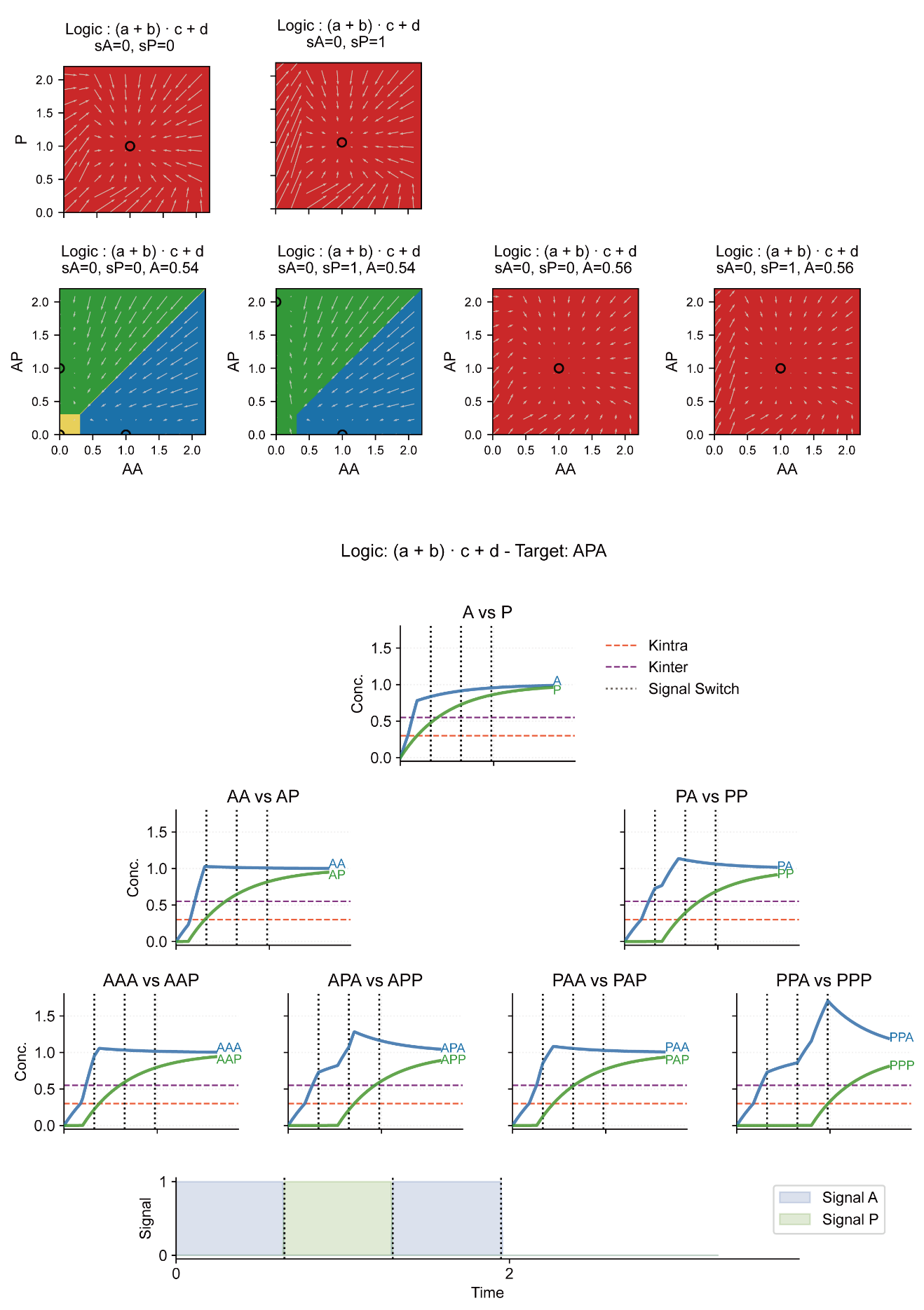

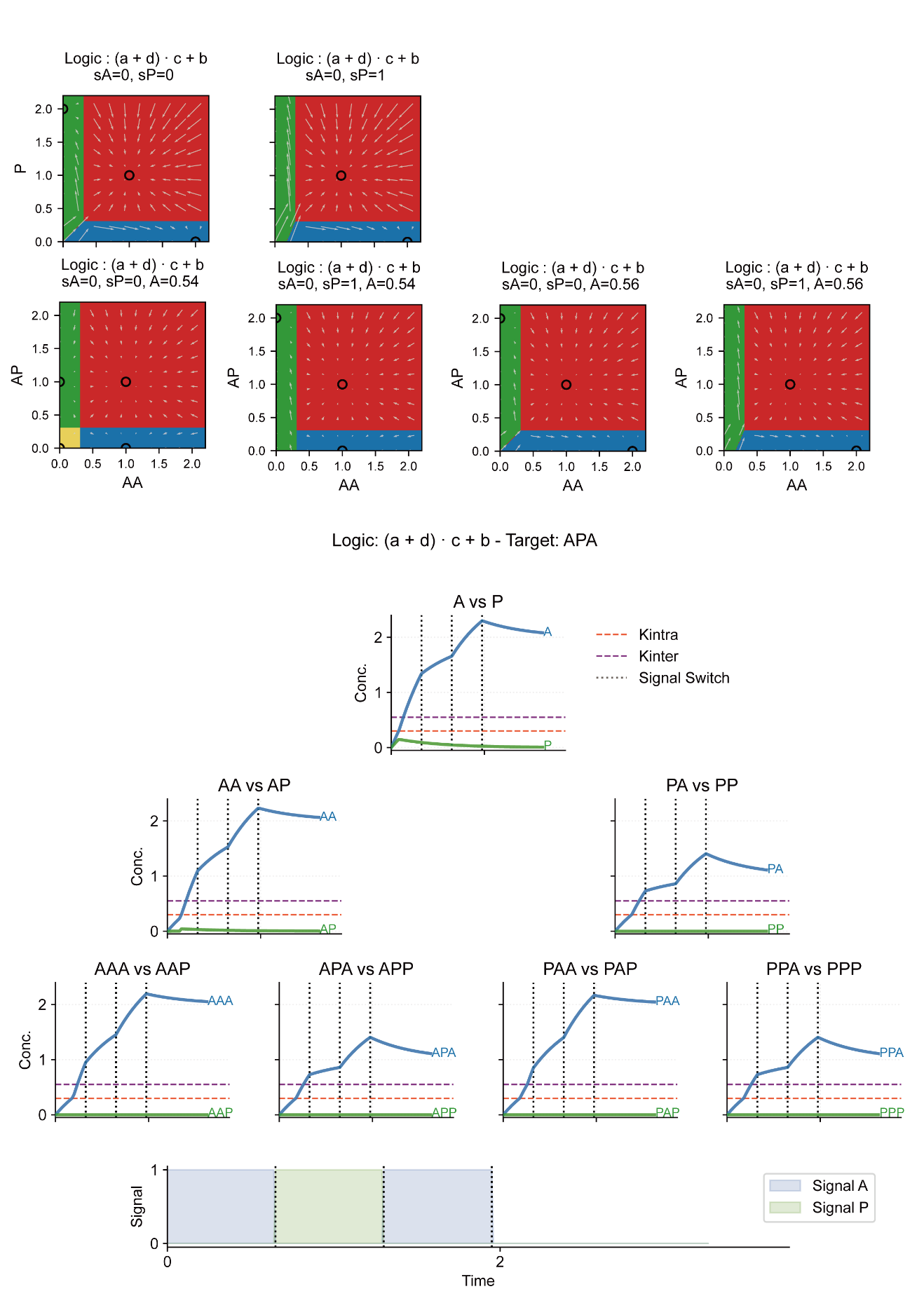

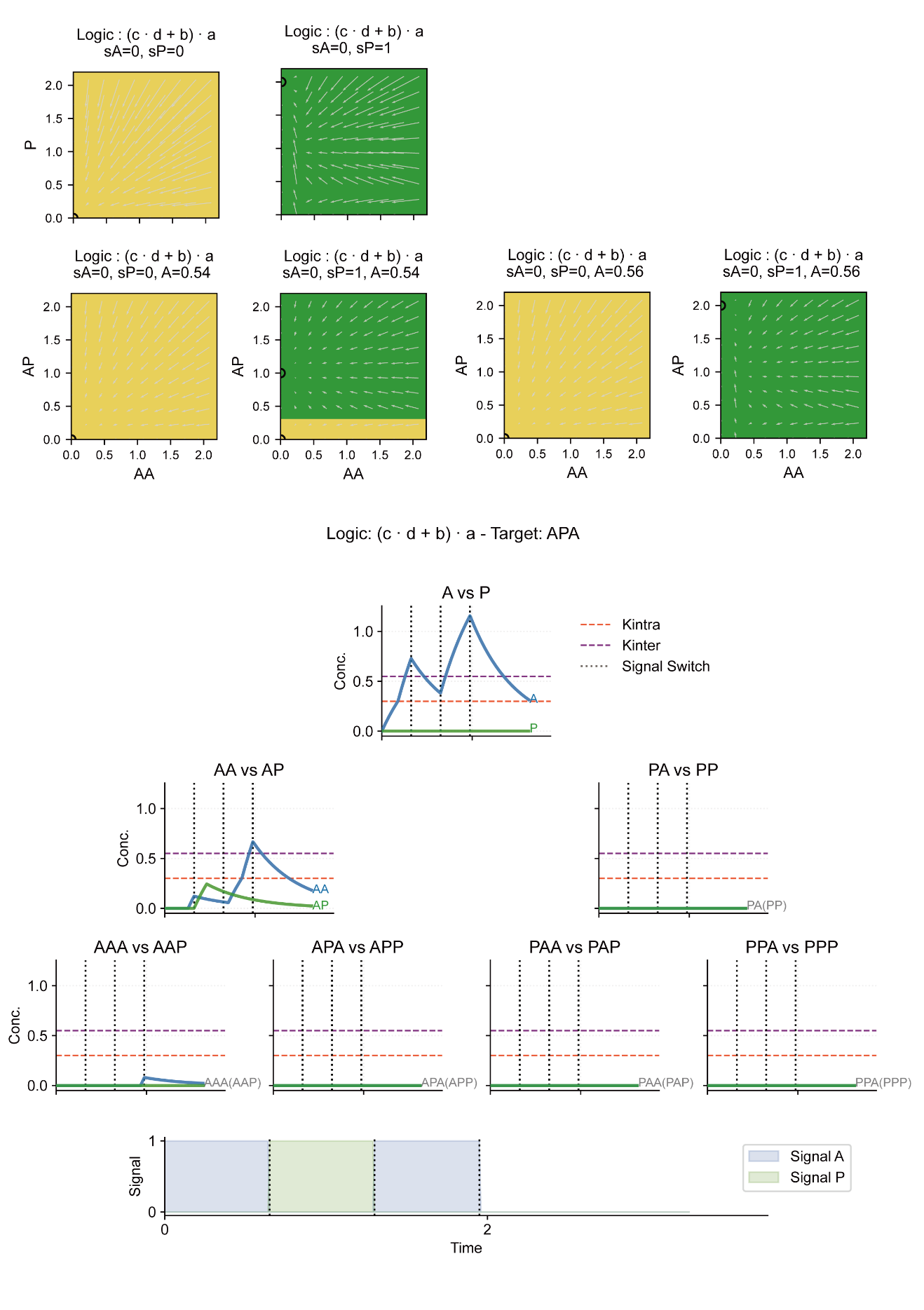

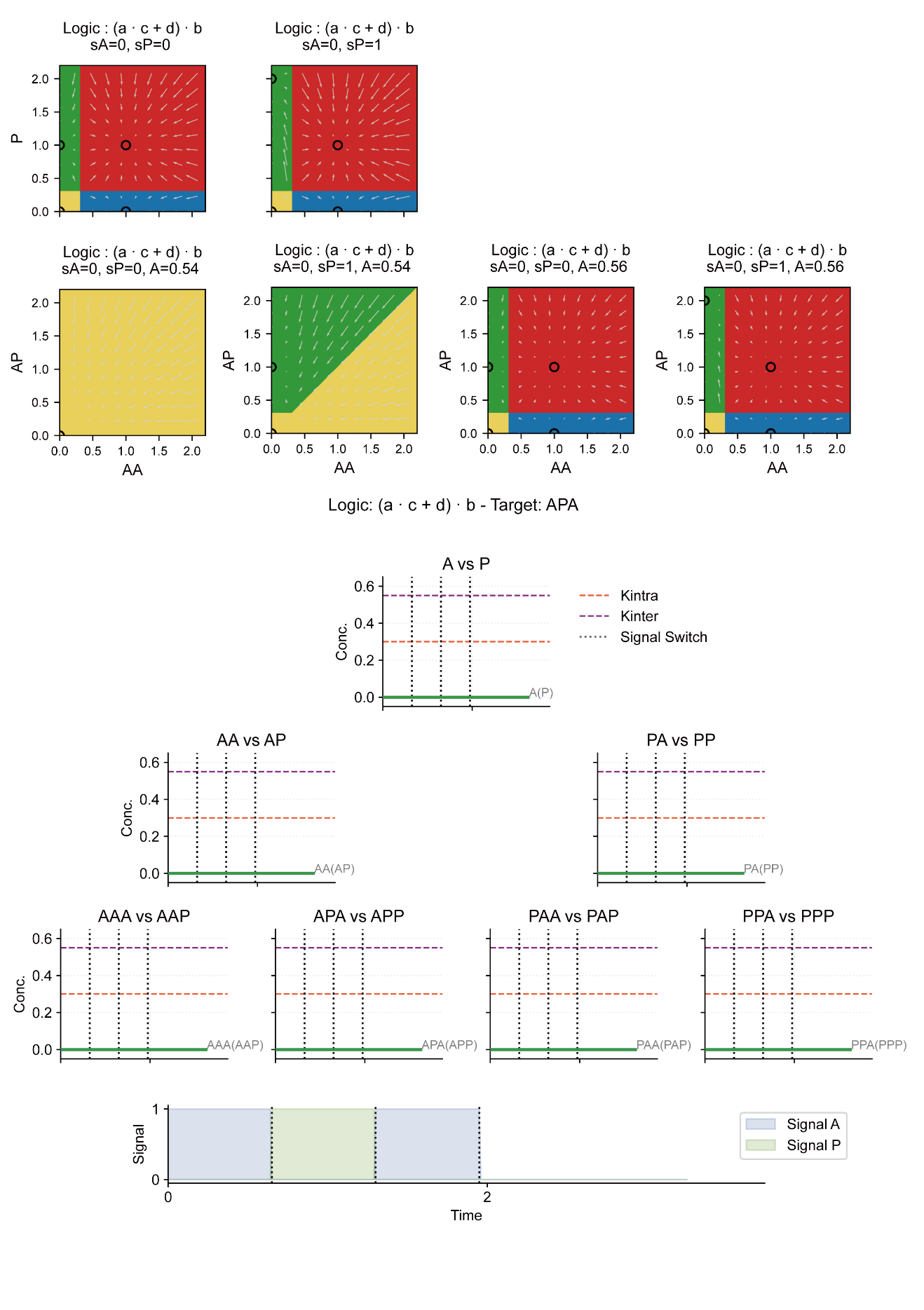

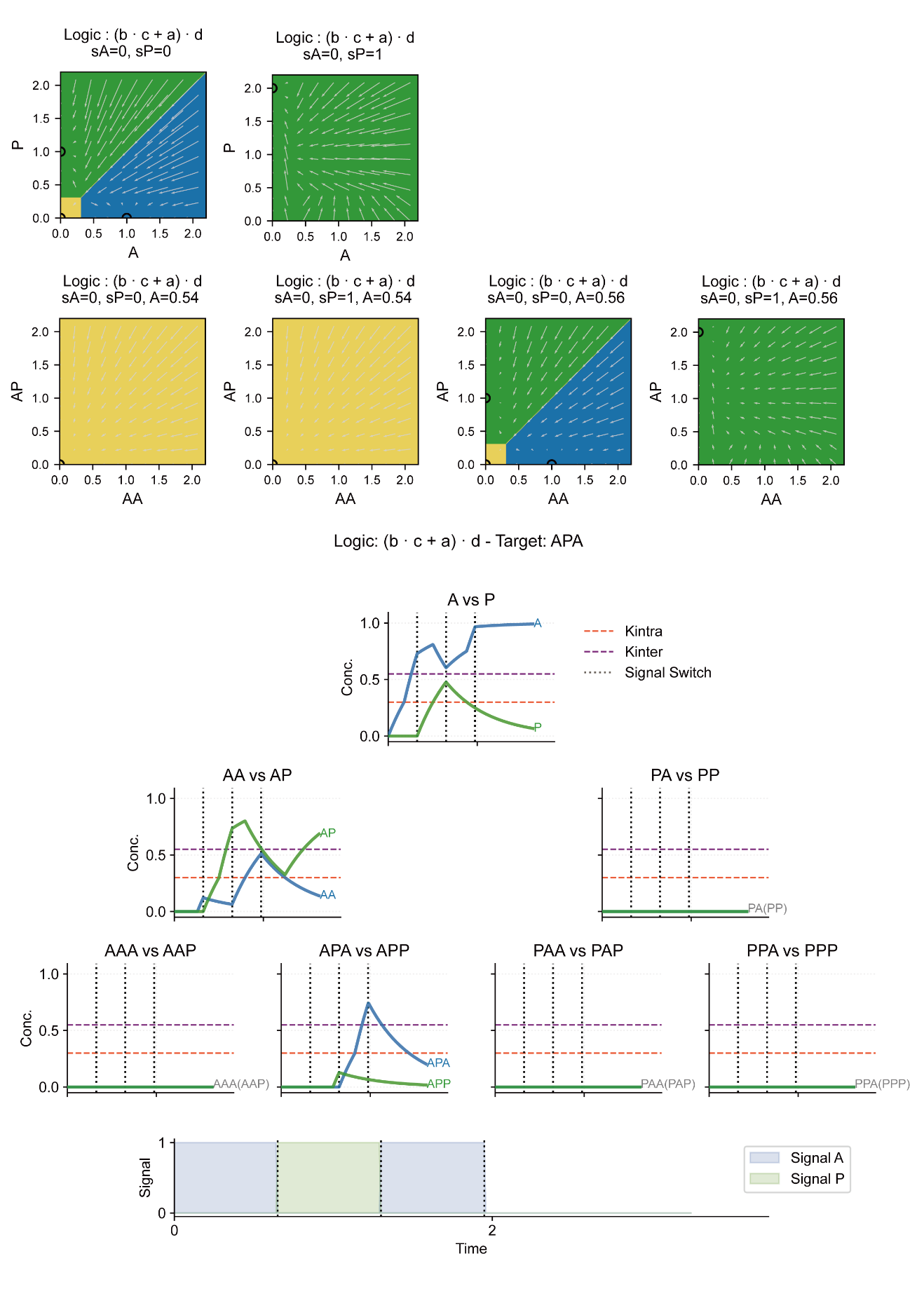

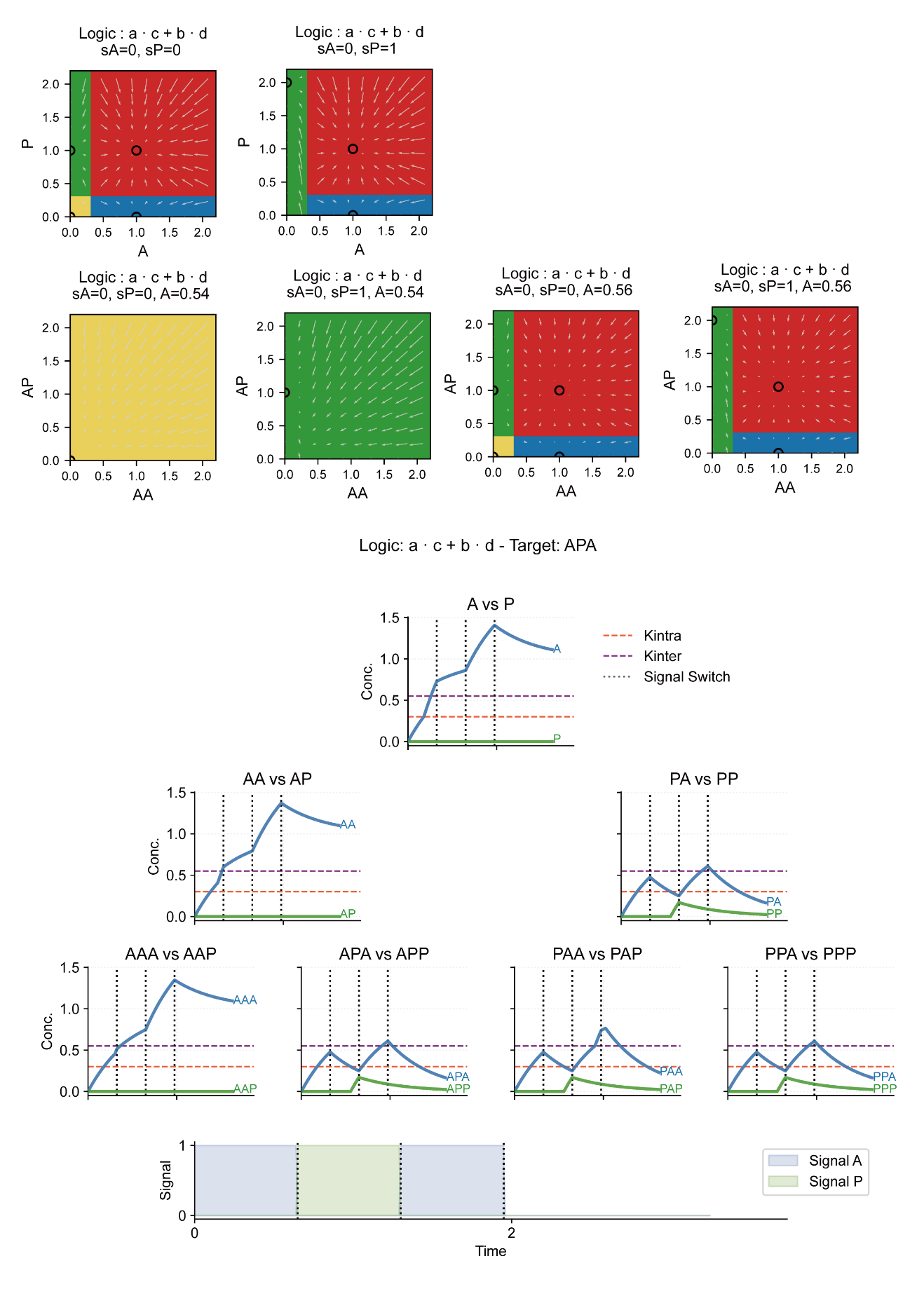

**Figure S5:** Clustering results of failure reasons.

The 52 combinatorial logics were systematically evaluated and clustered according to their specific topological defects observed in the first-layer (A-P) phase diagrams. Six binary criteria were utilized to characterize the failure modes: (1) Saddle Stem at S=(0,0): the double-low stem cell state acts as an unstable saddle point; (2) Only double-High at S=(0,0): the double-low stem cell state does not exist; (3) No Diff. State at S=(0,0): the absence of differentiated attractors prior to induction; (4) Stem Persists Under Induction: the abnormal persistence of the stem cell attractor despite signal induction; (5) No Diff. State Under Induction: the failure to generate differentiated states under inducing conditions; and (6) Inter-layer Decoupling: correct intra-layer regulation but failed inter-layer regulation. Dark red blocks in the heatmap indicate the presence of a specific error type for a given logic. Detailed evaluation metrics and categorizations for all logics are provided in Table S2.

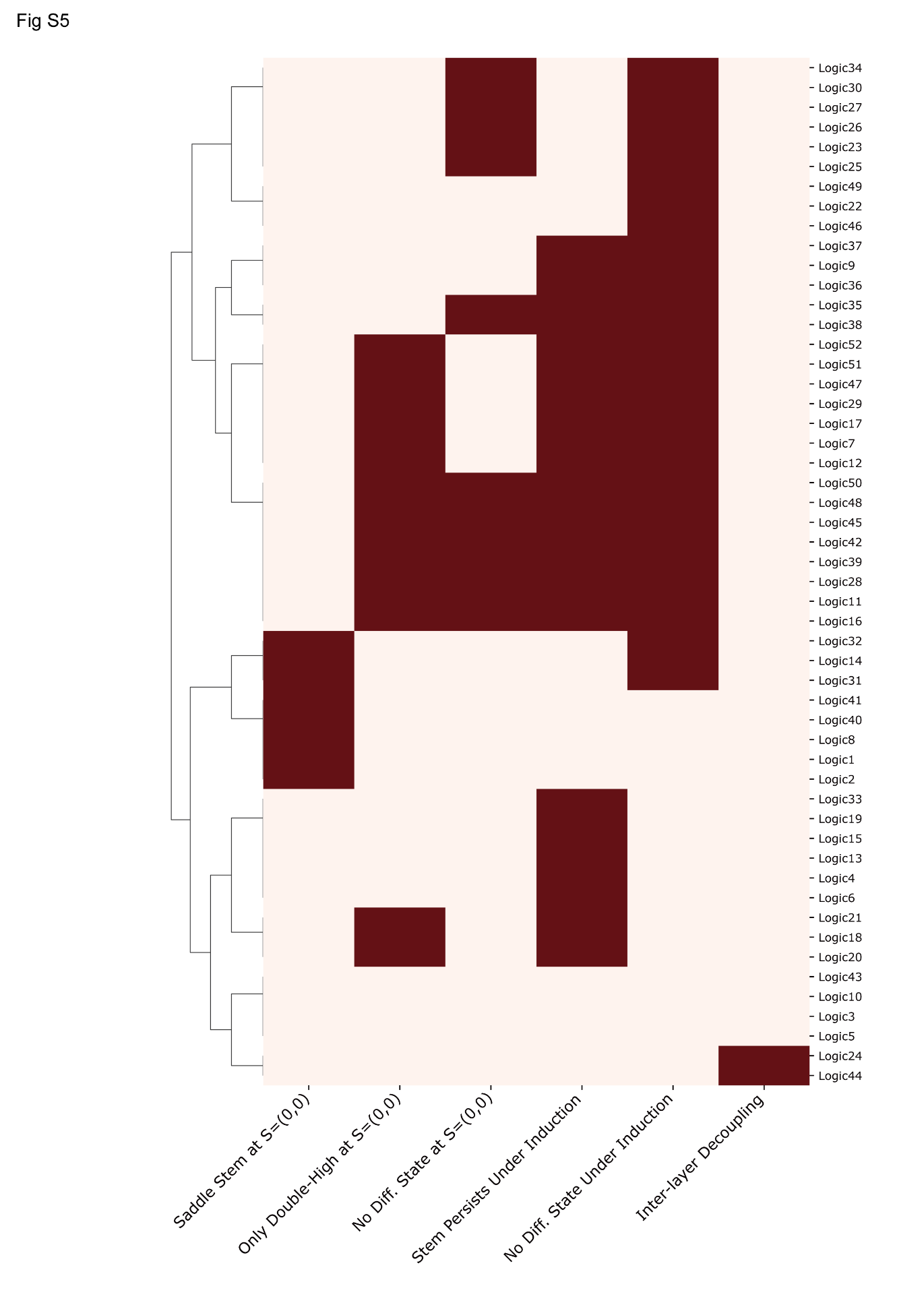

**Figure S6:** Numerical simulations showing the variation of $\langle\Delta t\rangle$ with Kintra and Kinter. (Related to Figure 3B).

Under the step-function approximation with a fixed blank interval ($\Delta t2=0$), ten representative target lineages were randomly sampled from the complete set of 64 possible targets of length L=6. For each parameter pair of Kintra and Kinter, we computationally scanned for the intersection of the permissible signal duration ($\Delta t$) windows that successfully induced all 10 sampled targets. The heatmap displays the average value of this overlapping $\Delta t$ window, denoted as $\langle\Delta t\rangle$. The color gradient from dark blue to yellow indicates increasing values of $\langle\Delta t\rangle$, whereas the white regions denote parameter regimes where no common successful $\Delta t$ window exists. The numerical results demonstrate that the required signal duration $\langle\Delta t\rangle$ monotonically increases as either Kintra or Kinter increases, which is in robust agreement with the analytical predictions presented in Figure 3B.

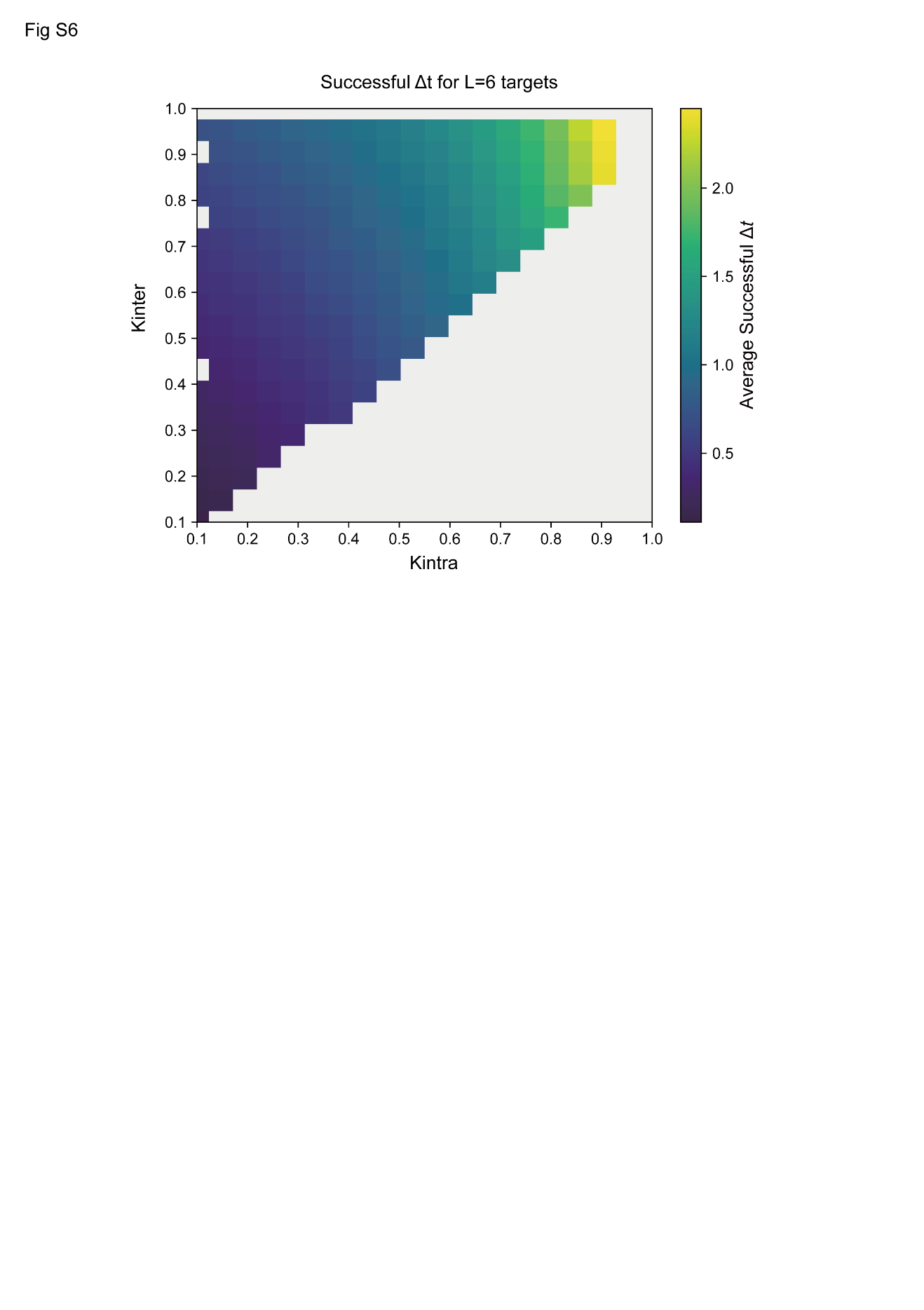

**Figure S7:** Dynamics simulations demonstrating the reset effect. (Related to Figure 3C).

Dynamic simulations of a representative logic configuration inducing three distinct target lineages (APAP, AAPA, and AAAP) are shown to illustrate the reset mechanism. Dark grey vertical dashed lines indicate the predefined time points of external signal switching. Light grey vertical dashed lines mark the moments when the current-layer transcription factor crosses the inter-layer threshold (Kinter), which initiates the expression of the downstream-layer factor.

During the induction of the APAP target (top panel), the temporal gap between the light and dark grey lines remains constant across layers. This indicates an absence of temporal error accumulation, owing to the reset effect provided by the alternating sA and sP signals.

For the AAPA target (middle panel), two consecutive sA pulses cause an initial error accumulation, evidenced by an increased distance between the second set of grey lines; however, the subsequent switch to sP triggers the reset effect, returning the third temporal gap to the baseline level.

Conversely, for the AAAP target (bottom panel), three consecutive sA pulses result in unmitigated, cascading error accumulation. This cumulative deviation ultimately causes the fourth-layer transcription factor to erroneously perceive the third-layer signal pulse, culminating in induction failure.

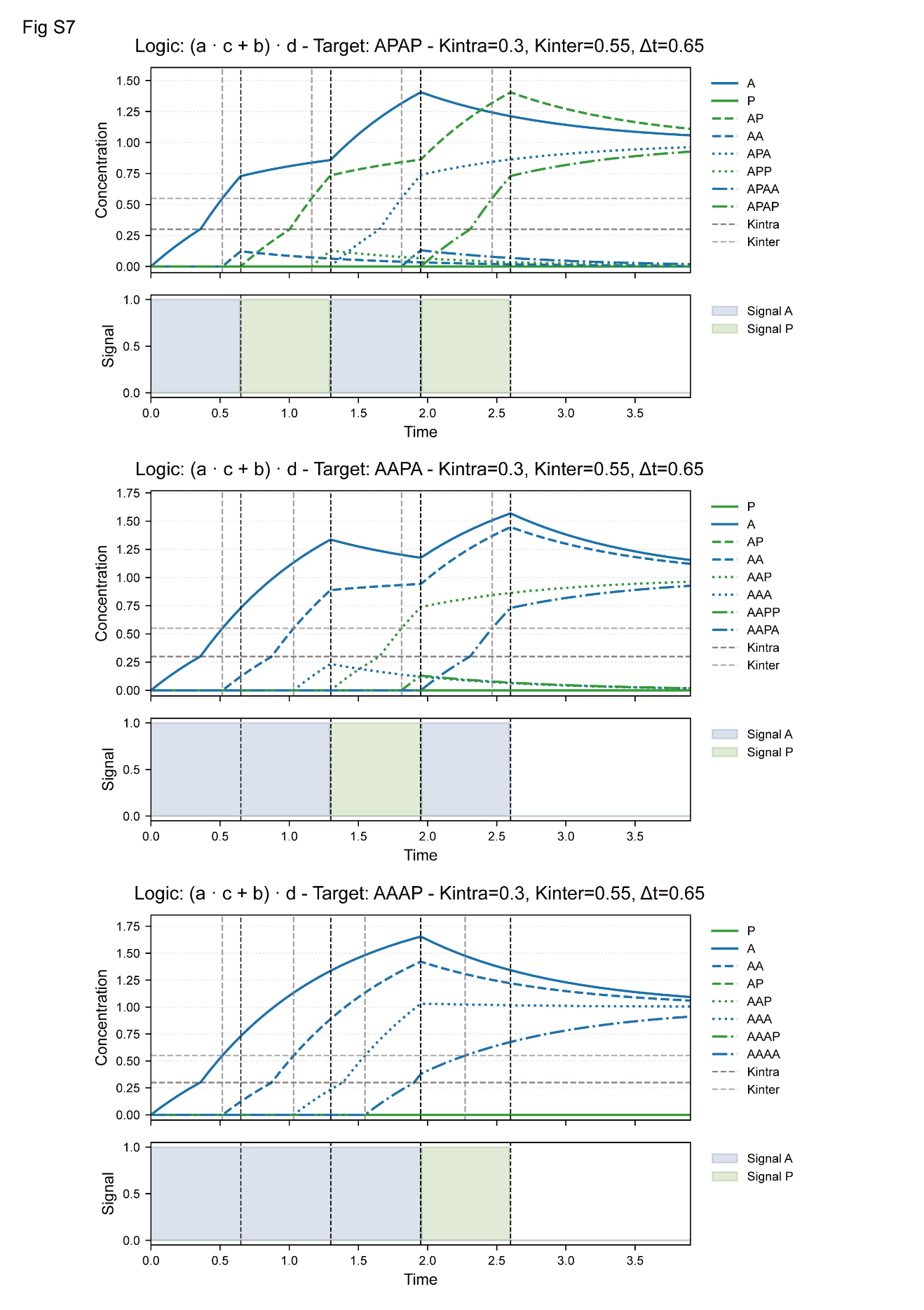

**Figure S8:** Effect of the non-induction interval *Δt2* on the permissible *Δt* window.

This figure illustrates the effect of extending the non-induction blank interval ($\Delta t2=0,0.5,1,2$) on the allowed signal duration ($\Delta t$) window for representative target lineages of length 6. The threshold parameters are fixed at Kintra = 0.3 and Kinter = 0.5. The y-axis displays the permissible range of $\Delta t$ that guarantees successful induction for each specific target sequence listed on the x-axis. The bars are color-coded according to the "Initial Run (y)" metric, which denotes the maximum length of the consecutive identical signals before a signal switching (e.g., repeated sA or sP inputs) before the first signal switch. Notably, as $\Delta t2$ increases, the allowed $\Delta t$ windows expand significantly, particularly for sequences with larger y values that are typically highly susceptible to error accumulation. This result demonstrates that introducing a blank interval effectively dissipates accumulated temporal errors, thereby relaxing the timing constraints and enhancing the overall robustness of the dynamic induction.

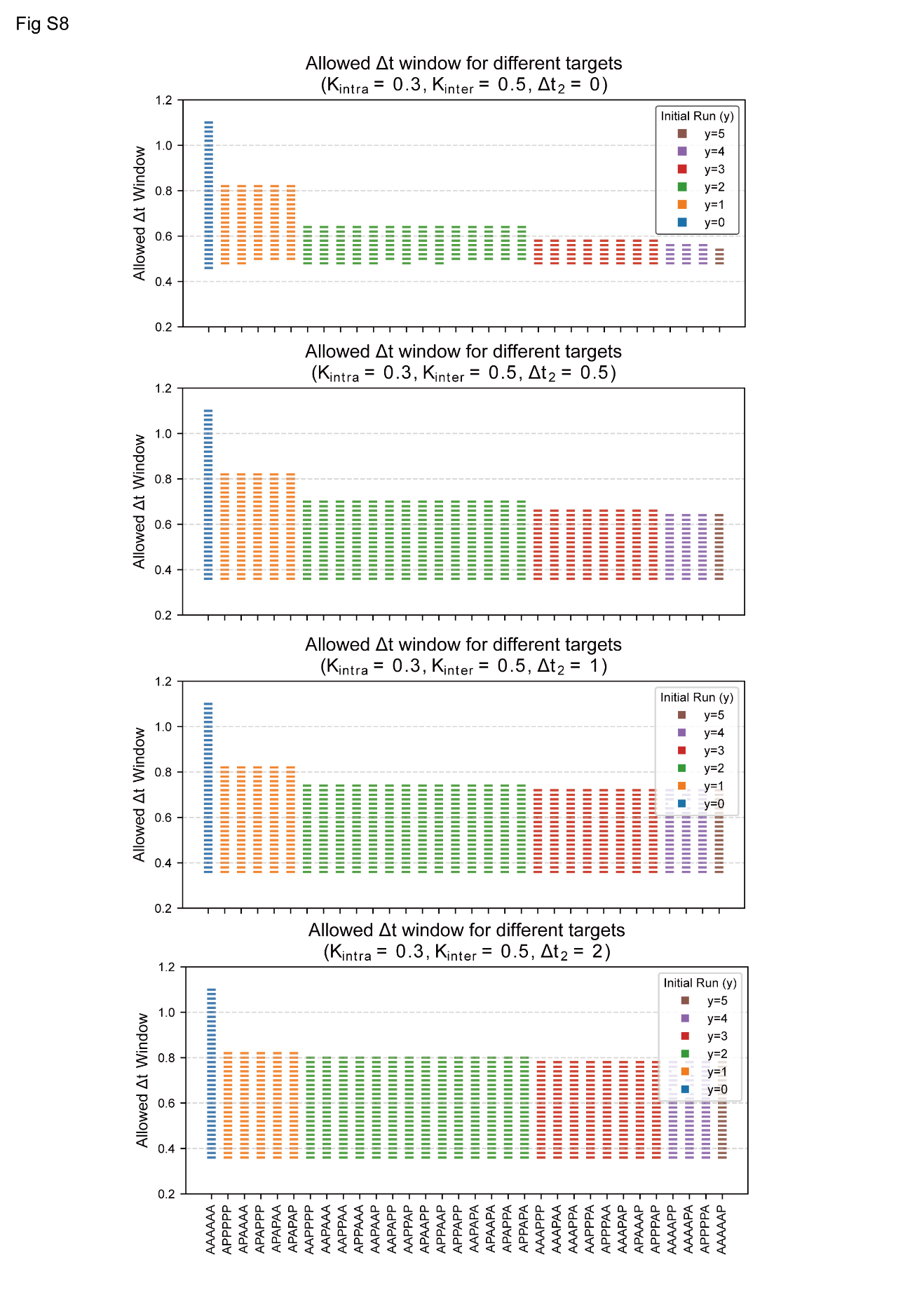

**Figure S9:** Dynamics simulations showing leaky expression under Hill functions.

(A) Dynamic simulation of the representative logic inducing the APA target using ideal Step functions. Under this approximation, downstream transcription factors remain completely inactive (concentration strictly at zero) until their upstream parent crosses the inter-layer threshold (Kinter).

(B) Dynamic simulation of the same logic and target using Hill functions. Unlike the Step function model, Hill functions intrinsically permit basal activation before thresholds are strictly met. The thick red bars on the x-axis highlight the specific periods of "leaky expression," during which downstream factors (e.g., the second-layer nodes) begin to accumulate prematurely before their upstream activator fully reaches Kinter.

In both panels, blue and green curves denote A-terminal and P-terminal transcription factors, respectively. The bottom subpanels display the alternating external signal sequence of sA and sP. Horizontal dashed lines indicate the Kintra and Kinter thresholds, while vertical dashed lines mark the temporal events of signal switching and threshold-crossing.

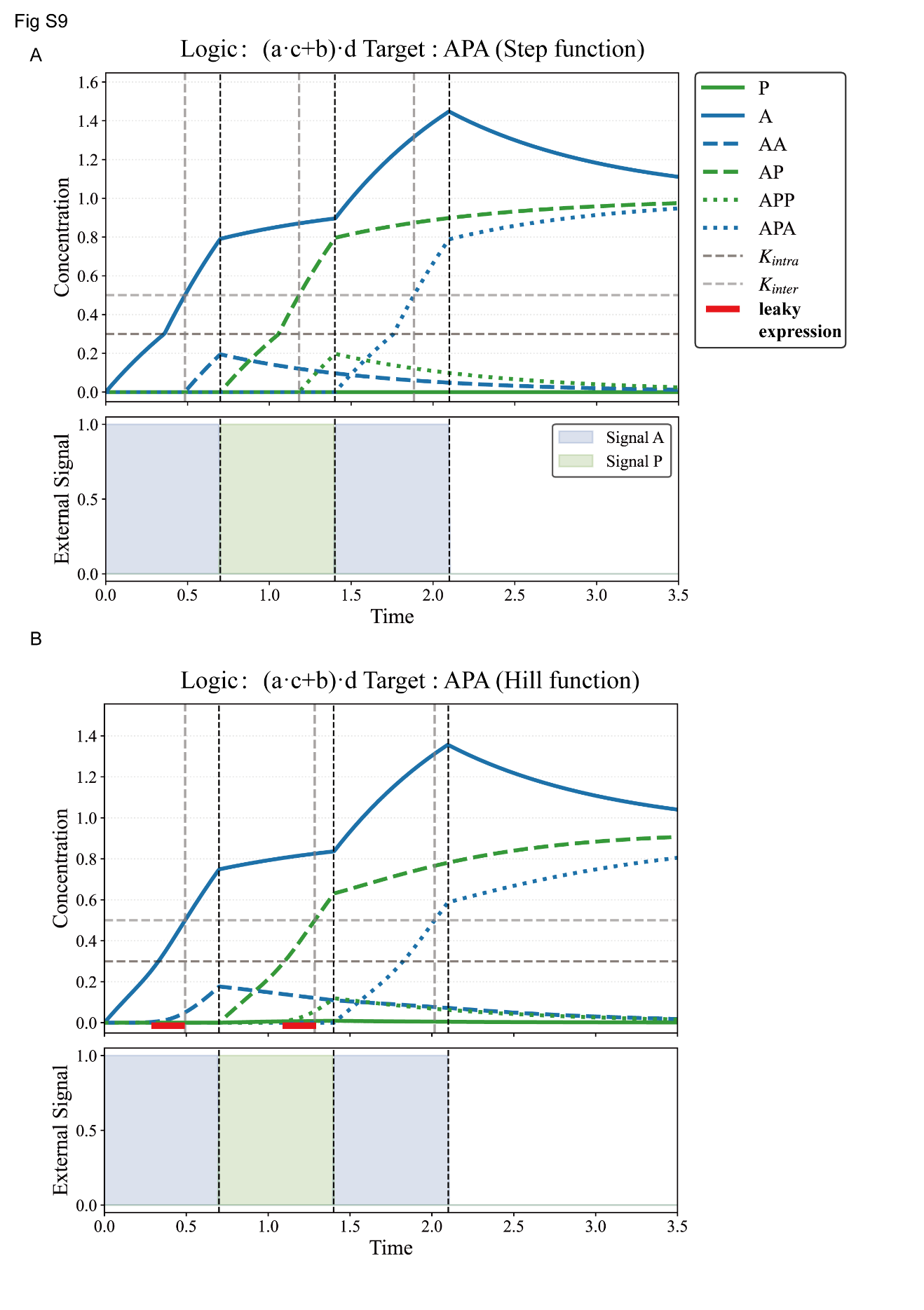

**Figure S10:** The dependence of permissible $K_{intra}$ on the Hill coefficient $n$ under Hill functions.

The y-axis denotes the maximum permissible Kintra, strictly defined as the critical upper threshold required to successfully maintain the proper topological structure of the first-layer (A-P) basins of attraction. The x-axis represents the Hill coefficient n, plotted on a logarithmic scale. The monotonically increasing curve reveals that a higher degree of regulatory cooperation (larger n, approaching an ideal step function) relaxes the constraint on Kintra, allowing the dynamic system to tolerate higher intra-layer thresholds. Conversely, for systems with smaller n values (representing more continuous and less switch-like responses), Kintra must remain tightly restricted to a lower regime to preserve the correct stable cell states and avoid topological failure.

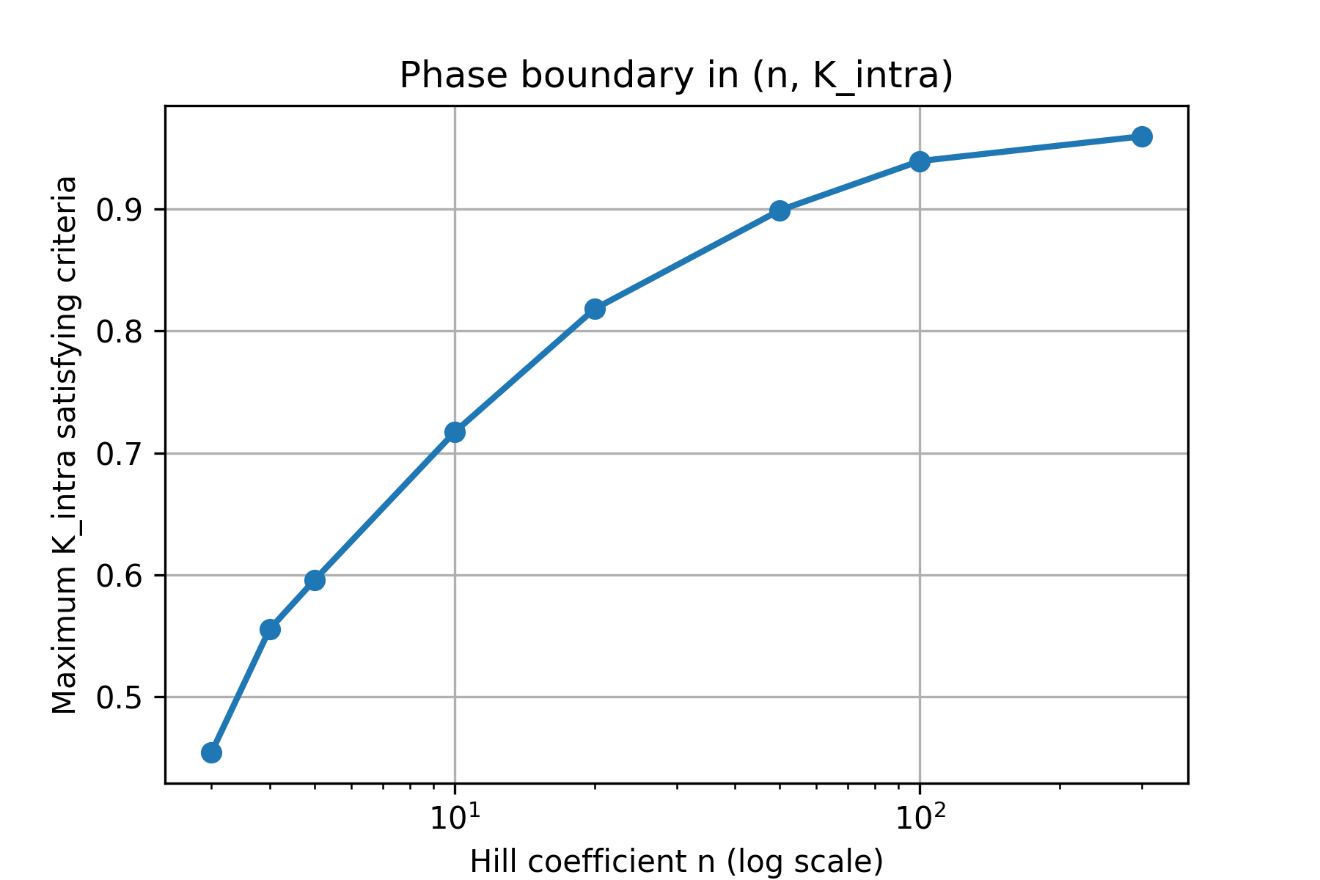

Figure S11: Workflow and representative results of the high-throughput parameter screening under the generalized dynamical framework. (Related to Note S4).

Flowchart detailing the high-throughput screening pipeline. The process begins with Latin Hypercube Sampling (LHS) to generate 16,000 system models, followed by a rigorous topological evaluation of the static attraction basins for both Layer 1 and Layer 2. A single-step micro-perturbation is artificially introduced to eliminate pseudo-attractor misclassifications at saddle points. Finally, dynamic simulations are executed from a pre-equilibrated genuine low-stem state at S=0.5, scanning for successful signal duration dt windows under a fixed blank gap $\Delta t2=10$.

Static phase diagrams for a representative successful parameter set (Row 4062, Logic (c+d)*b) that passed the stringent topological screening. The top row illustrates the first-layer (A-P) phase diagrams under S=0, 0.5, and 1, demonstrating the required multistability at the basal signal (S=0.5) and the bifurcation of the stem cell attractor at S=0 or 1. The subsequent three rows display the second-layer (AA-AP) phase diagrams evaluated at 9 fine-grained conditions, capturing the cascading effects when the upstream node A is at the stem, fully induced (A_high), or dynamically shrinking (A_shrink) states.

Dynamic simulations of the same successful parameter set, showing the successful multi-stage induction of the AAAP (left subpanel) and APAP (right subpanel) target sequences. The trajectories utilize the identified permissible signal window dt=10.4 and the fixed non-induction interval $\Delta t2=10$, conclusively validating that the specific static basin morphology is a strong predictive hallmark for dynamic induction success.
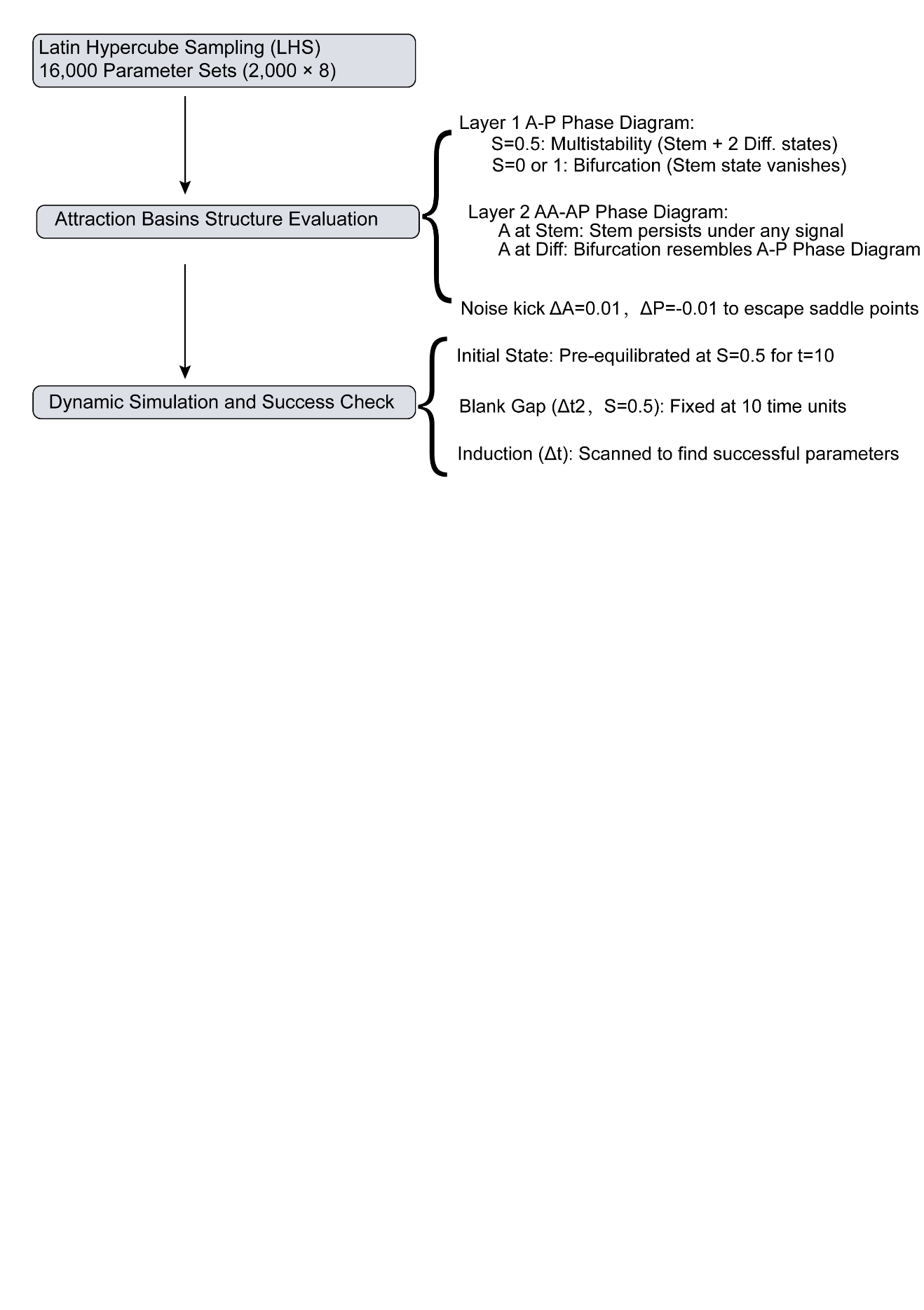

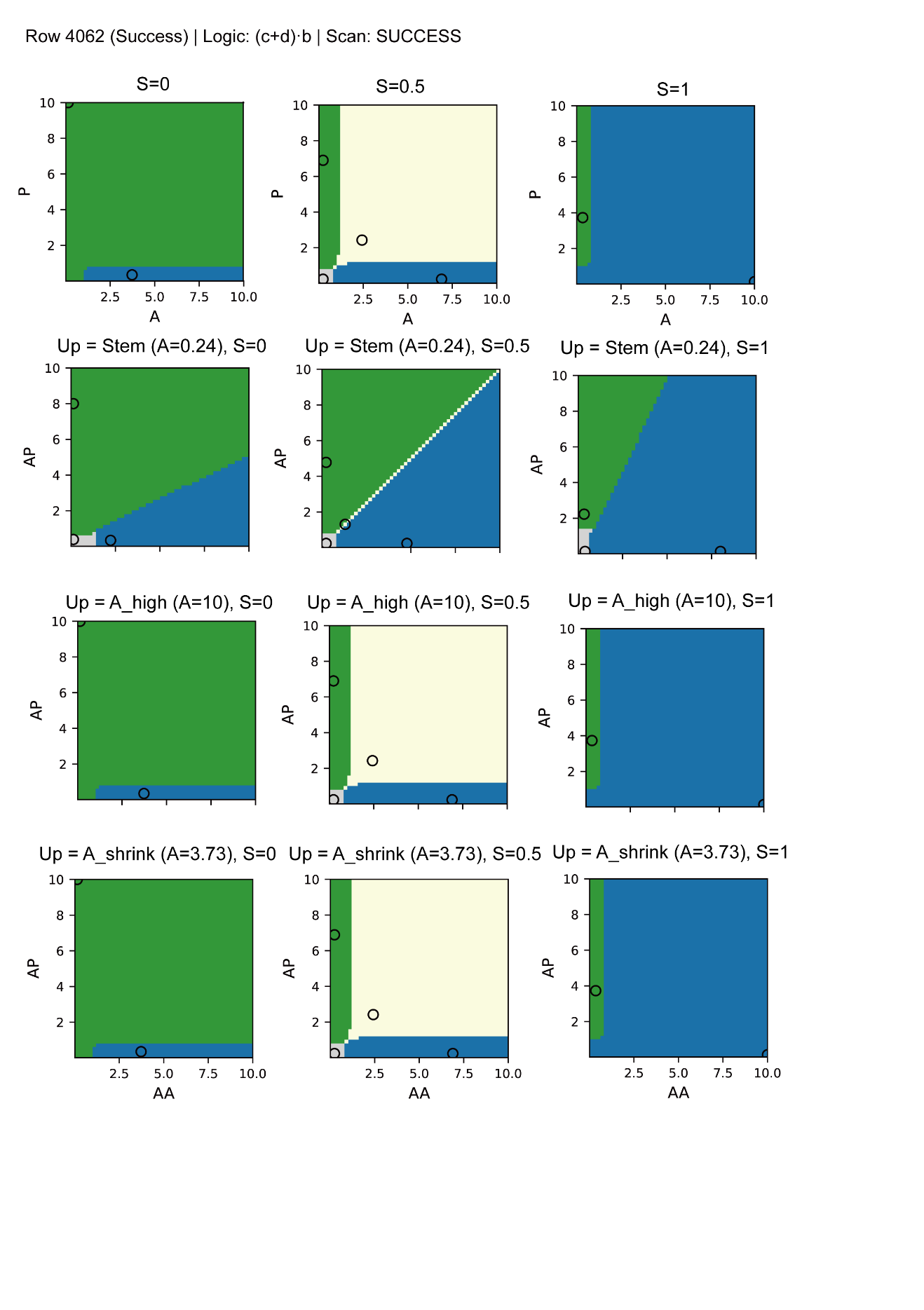

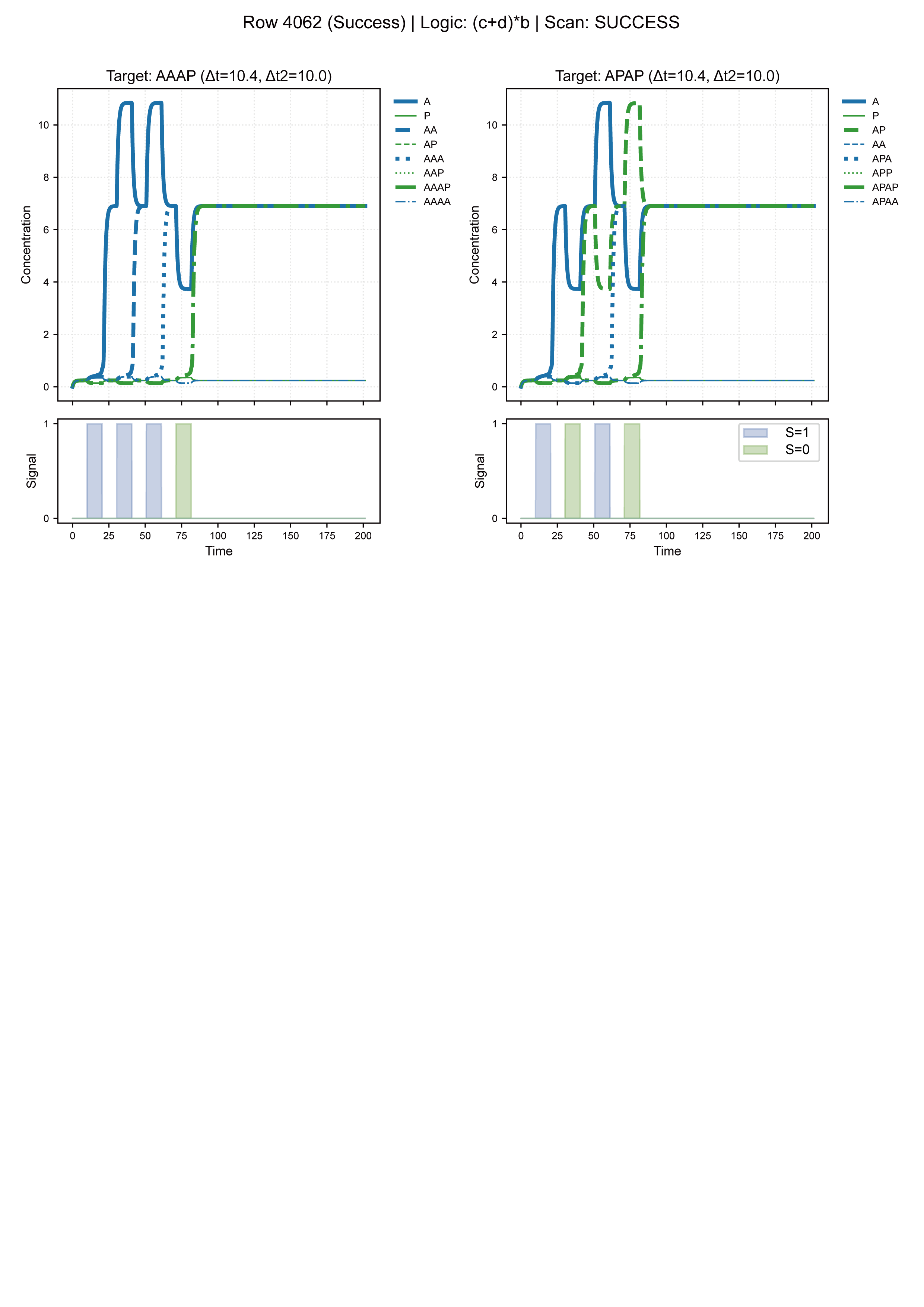
